# ipaA triggers vinculin oligomerization to strengthen cell adhesion during *Shigella* invasion

**DOI:** 10.1101/773283

**Authors:** Cesar Valencia-Gallardo, Daniel-Isui Aguilar-Salvador, Hamed Khakzad, Charles Bou-Nader, Christian Malosse, Diogo Borges Lima, Chakir Bello, Benjamin Cocom-Chan, Nicole Quenech’Du, Bilal Mazhar, Delphine Javelaud, Jacques Fattaccioli, Alain Mauviel, Marc Fontecave, Atef Asnacios, Julia Chamot-Rooke, Lars Malmström, Guy Tran Van Nhieu

**Affiliations:** Equipe Communication Intercellulaire et Infections Microbiennes, Centre de Recherche Interdisciplinaire en Biologie (CIRB), Collège de France, 75005 Paris, France; Institut National de la Santé et de la Recherche Médicale U1050, 75005 Paris, France; Centre National de la Recherche Scientifique UMR7241, 75005 Paris, France; MEMOLIFE Laboratory of excellence and Paris Science Lettre; Institute for Computational Science, University of Zürich, Zürich, Switzerland; Swiss Institute of Bioinformatics, Lausanne, Switzerland; Laboratoire de Chimie des Processus Biologiques, Collège De France, 75005 Paris, France; Centre National de la Recherche Scientifique UMR8229, 75005 Paris, France; Mass Spectrometry for Biology Utechs, Institut Pasteur, USR 2000, CNRS, 75015 Paris, France; Institut Curie, PSL Research University, INSERM U1021, CNRS UMR3347, Team “TGF-ß and Oncogenesis“, Equipe Labellisée LIGUE 2016, F-91400, Orsay, France; Université Paris-Sud, F-91400, Orsay, France; Centre National de la Recherche Scientifique UMR 3347, 91400 Orsay, France; PASTEUR, Département de Chimie, École Normale Supérieure, PSL University, Sorbonne Université, CNRS, 75005 Paris, France; Institut Pierre-Gilles de Gennes pour la Microfluidique, 75005 Paris, France; Laboratoire Matière et Systèmes Complexes, UMR 7057 CNRS & Université Paris Diderot, Sorbonne Paris Cité, Paris, France; Division of Infection Medicine, Department of Clinical Sciences, Lund University, Lund, Sweden

**Keywords:** Shigella, invasion, vinculin, adhesion

## Abstract

The *Shigella* effector IpaA co-opts the focal adhesion protein vinculin to promote bacterial invasion. Here, we show that IpaA triggers an unreported mode of vinculin activation through the cooperative binding of its three vinculin-binding sites (VBSs) leading to vinculin oligomerization via its D1 and D2 head subdomains and highly stable adhesions resisting actin relaxing drugs. Using cross-linking mass spectrometry, we found that while IpaA VBSs1-2 bound to D1, IpaA VBS3 interacted with D2, a subdomain masked to other known VBSs. Structural modeling indicated that as opposed to canonical activation linked to interaction with D1, these combined VBSs interactions triggered major allosteric changes leading to D1D2 oligomerization. A cysteine-clamp preventing these changes and D1D2 oligomerization impaired growth of vinculin microclusters and cell adhesion. We propose that D1D2-mediated vinculin oligomerization occurs during the maturation of adhesion structures to enable the scaffolding of high-order vinculin complexes, and is triggered by *Shigella* IpaA to promote bacterial invasion in the absence of mechanotransduction.

**Summary:** The *Shigella* IpaA effector binds to cryptic vinculin sites leading to oligomerization via its head domain. This vinculin oligomerization mode appears required for the maturation and strengthening of cell adhesion but is co-opted by invasive bacteria independent of actomyosin contractility.

## Introduction

*Shigella*, the causative agent of bacillary dysentery, invades epithelial cells by injecting type III effectors that locally reorganize the actin cytoskeleton (Ogawa, Handa et al. 2008, Valencia-Gallardo, Carayol et al. 2015, Mattock and Blocker 2017). *Shigella* invasion involves limited contacts with host cells and critically depends on the type III effector IpaA that promotes cytoskeletal anchorage by targeting the focal adhesion proteins talin, and vinculin (Romero, Grompone et al. 2011, Valencia-Gallardo, Carayol et al. 2015, Valencia-Gallardo, Bou-Nader et al. 2019). During integrin-mediated cell adhesion, talin acts as a mechanosensor by exposing its vinculin binding sites (VBSs) that recruit and activate vinculin, reinforcing anchorage to the actin cytoskeleton in response to mechanical load (Humphries, Wang et al. 2007, Parsons, Horwitz et al. 2010, Lavelin, Wolfenson et al. 2013). *Shigella* cannot generate the type of mechanical load required for strong cytoskeletal anchorage, therefore, scaffolding of talin, and vinculin at bacterial invasion sites exclusively relies on IpaA. IpaA contains three vinculin binding sites (VBSs) in its carboxyterminal moiety, with diverse functions inferred from the crystal structures of complexes containing the VBS peptide (Izard, Tran Van Nhieu et al. 2006, Tran Van Nhieu and Izard 2007). Vinculin is classically described as a head domain (Vh) connected to a tail domain (Vt) by a flexible linker (Bakolitsa, Cohen et al. 2004). Vh contains three repetitions (D1-D3) of a conserved domain consisting of two antiparallel α-helix-bundles and a fourth α-helix bundle D4 (Bakolitsa, Cohen et al. 2004). A proline-rich unstructured linker bridges D4 and a five-helix bundle Vt containing the carboxyterminal F-actin binding domain (Bakolitsa, Cohen et al. 2004). Under its inactive folded form, intramolecular interactions between Vh and Vt prevent ligand binding. IpaA VBS1, as for all VBSs described to activate vinculin, interacts with the first helical bundle of the D1 domain, promoting major conformational changes that disrupt the Vh-Vt intramolecular interactions and free the vinculin F-actin binding region (Izard, Evans et al. 2004). IpaA VBS2, in contrast, interacts with the second helical bundle of D1 (Tran Van Nhieu and Izard 2007), hence, its association with IpaA VBS1 results in a very high affinity and stable IpaA VBS1-2:D1 complex, with an estimated K_D_ in the femtoM range (Tran Van Nhieu and Izard 2007). Functional evidence indicates that IpaA VBS3 cooperates with IpaA VBS1-2 to stimulate bacterial invasion (Park, Valencia-Gallardo et al. 2011, Valencia-Gallardo, Bou-Nader et al. 2019). IpaA VBS3, as an isolated peptide, acts as IpaA VBS1 by interacting with the vinculin D1 first helical bundle and promotes vinculin activation (Park, Valencia-Gallardo et al. 2011). IpaA VBS3, however, can also interact with talin to stimulate bacterial capture by filopodia during the early invasion phase (Valencia-Gallardo, Bou-Nader et al. 2019). The structural data indicate that IpaA VBS3 stabilizes the H1-H4 helix bundle expected to form in a partially stretched talin conformer at the low force range exerted by filopodia (Valencia-Gallardo, Bou-Nader et al. 2019). Intriguingly, IpaA VBS3 shares with talin VBS10 (H46) the ability to bind to vinculin and talin H1H4, suggesting a complex interplay between talin and vinculin during mechanotransduction (Valencia-Gallardo, Bou-Nader et al. 2019). Unlike talin VBSs, IpaA VBSs are not buried into helix bundles, presumably enabling targeting of vinculin and talin in a serendipitous manner. This property allows the targeting of talin adhesions by IpaA VBS3 during filopodial capture, but also questions the role of its vinculin binding activity with respect to that of IpaA VBS1, 2 during adhesion to the cell cortex, presumably at higher force ranges that are incompatible with IpaA VBS3 binding to talin.

In addition to strengthening cytoskeletal anchorage, vinculin has also been implicated in the bundling of actin filaments through dimerization via its tail domain (Vt), triggered by F-actin or phosphatidylinositol(4, 5) bisphosphate (PIP2) binding (Johnson and Craig 2000, Janssen, Kim et al. 2006, Chinthalapudi, Rangarajan et al. 2014). Consistent with a key role in Vt-mediated actin bundling, mutations in Vt that prevent PIP2-binding lead to defects in focal adhesion dynamics and formation (Chinthalapudi, Rangarajan et al. 2014). Also, mutations that prevent Vt dimerization or alter C-terminal hairpin involved in actin bundling lead to defects in focal adhesion formation and cell spreading, although the correlation between F-actin bundling activity and the amplitude of adhesion defects is unclear (Shen, Tolbert et al. 2011). Of interest, upon activation, vinculin is known to promote the scaffolding of adhesion components during focal adhesion growth and maturation, a process that may also implicate its oligomerization (Thompson, Tolbert et al. 2013). The role of Vt-induced dimerization in scaffolding, however, remains unclear since it cannot simply explain the formation of high-order complexes. The formation of these latters could implicate the recruitment of other vinculin-binding partners or vinculin oligomerization mechanisms other than through Vt, possibly through Vh-Vh interactions observed in the so-called “parachute” structures (Molony and Burridge 1985). Here, we investigated the role of vinculin at the cell cortical sites of *Shigella* invasion, where fully activated talin is not expected to bind to IpaA VBS3 and all three IpaA VBSs are expected to bind target vinculin (Valencia-Gallardo, Bou-Nader et al. 2019).

## Results

We first analyzed vinculin structures during cell challenge with *Shigella* strains expressing IpaA VBS3 derivatives deficient for talin-binding. As shown in Fig. 1a and as previously reported, *Shigella* triggers the IpaA-dependent recruitment of vinculin at phagocytic cups, as well as the formation of vinculin adhesion structures at the cell basal surface during cell invasion (Figs. 1a and 1b, WT). Vinculin recruitment at bacterial contact sites, however, was strongly reduced at actin foci induced by an *ipaA* mutant expressing IpaA VBS3 but not IpaA VBS1, 2 (Figs.1 a and 1b, VBS3; Fig. S1). For this strain, small vinculin patches were observed in instances at the immediate bacterial vicinity, but phagocytic cups surrounding invading observed for the wild-type strain were not detected (Figs. 1a, b, arrows). Complementation with IpaA VBS3 derivatives containing amino acid substitutions that affect talin-but not vinculin-binding gave similar results, consistent with a direct role of IpaA VBS3 in the recruitment of these vinculin residual levels (Figs. 1a and 1b, A495C and K498E; Fig. S1). Strikingly, however, IpaA VBS3 clearly stimulated the formation of large vinculin adhesion structures in the absence of IpaA VBS1, 2 (Figs. 1c, 1d and S1). As opposed to what was previously observed for talin-containing adhesions (Valencia-Gallardo, Bou-Nader et al. 2019), these large vinculin adhesions were induced to a similar extent with talin-binding deficient IpaA VBS3 derivatives.

**Figure 1.**
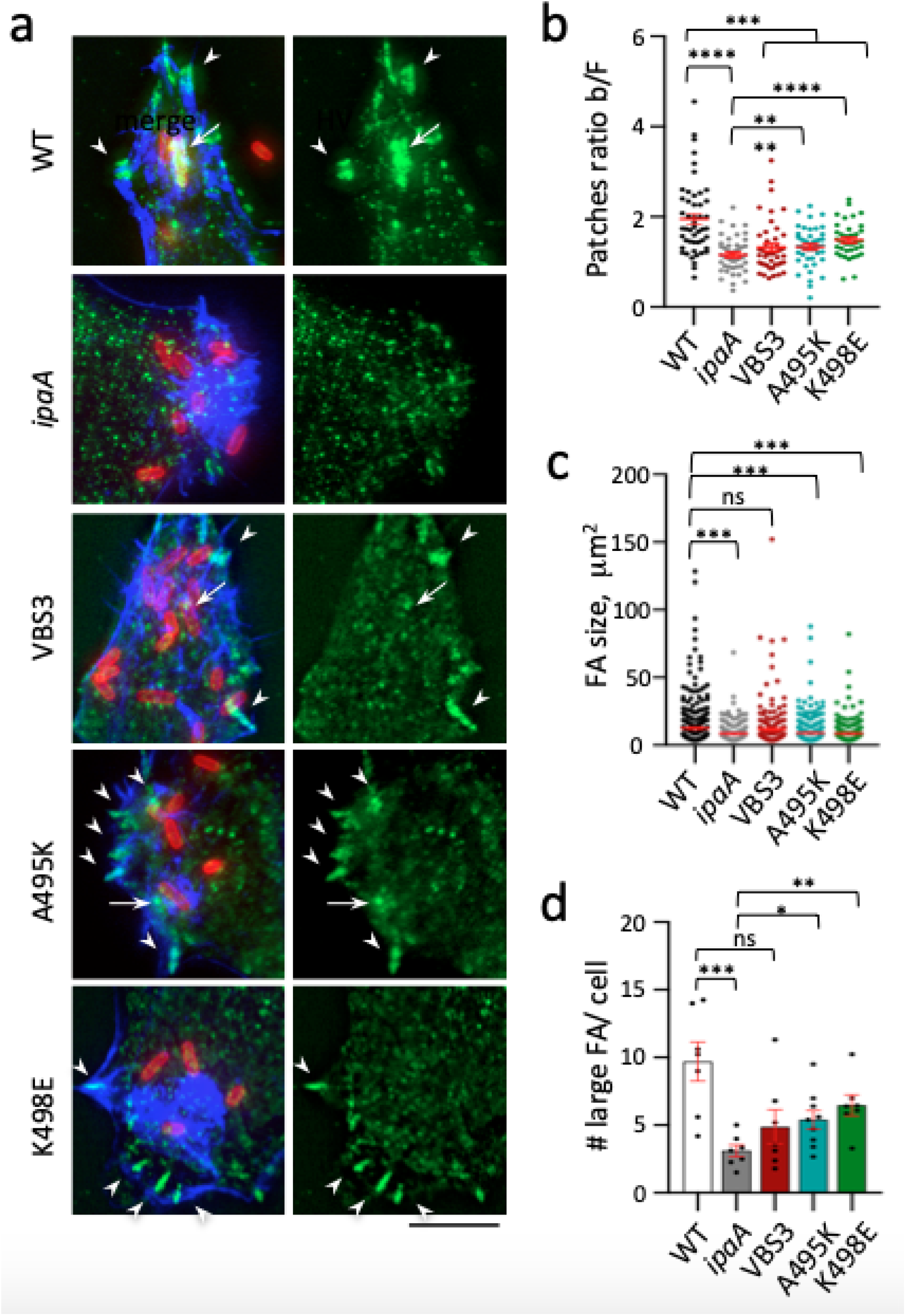
Vinculin recruitment at *Shigella* contact sites and distal adhesion structures during bacterial invasion. HeLa cells were challenged with bacteria for 30 min at 37°C, fixed and processed for immunofluorescence staining. *ipaA* mutant complemented with: full-length IpaA (WT); control vector (*ipaA*); IpaA ΔVBS1-2 (VBS3); IpaAΔVBS1-2 A495K (A495K); IpaAΔVBS1-2 K498E (K498E). **a**, representative micrographs. Merge: maximum projection of deconvolved confocal planes. HV: confocal plane corresponding to the cell basal surface. Red: bacteria; green: vinculin. Blue: F-actin. Vinculin recruitment at bacterial contact sites (arrows) and distal adhesion structures (arrowheads). Scale bar = 5 μm. **b**, vinculin recruitment at bacterial contact sites was quantified as the ratio of average fluorescence intensity of vinculin labeling associated with the bacterial body over that of actin foci (Materials and Methods, Fig. S1). The average ratio ± SEM is indicated. WT: 1.94 ± 0.11 (48 foci, N = 2); *ipaA*: 1.15 ± 0.05 (46 foci, N = 2); VBS3: 1.32± 0.08 (45 foci, N = 2); A495K: 1.33 ± 0.07 (42 foci, N = 2); K498E: 1.48 ± 0.06 (38 foci, N = 2). **c, d,** large vinculin adhesion structures were scored as detailed in the Materials and Methods section. **c,** average FA size ± SEM μm^2^: WT: 12.54 ± 0.76 (393 FAs, N = 2); *ipaA*: 8.48 ± 0.49 (201 FAs, N = 2); VBS3: 11.59 ± 1.08 (207 FAs, N = 2); A495K: 8.96 ± 0.45 (376 FAs, N = 2); K498E: 8.56 ± 0.44 (291 FAs, N = 2). **d,** average FA number per cell ± SEM: WT: 9.7 ± 1.44 (46 cells, N = 2); *ipaA*: 3.11 ± 0.43 (72 cells, N = 2); VBS3: 5.18 ± 1.43 (52 cells, N = 2); A495K: 5.4 ± 0.73 (78 cells, N = 2); K498E: 6.48 ± 0.77 (50 cells, N = 2). Mann and Whitney test: *: p < 0.05; **: p <0.01; ***: p <0.005; ****: p <0.001.

These results suggest that IpaAVBS3 binds to vinculin and argue for different but concerted roles of IpaA VBS1, 2 and IpaA VBS3 in the recruitment of vinculin at phagocytic cups or distal adhesion structures during *Shigella* invasion.

Previous analytical size exclusion chromatography (SEC) studies suggested that IpaA can bind to multiple vinculin molecules through its three VBSs (Park, Valencia-Gallardo et al. 2011). In the proposed model and akin to the model proposed for talin VBSs during mechanotransduction, each IpaA VBS binds to one vinculin molecule through interaction via the first bundle of the vinculin D1 subdomain, leading to its activation (Fig. 2a) (Park, Valencia-Gallardo et al. 2011). This view supports a redundant but not differential role for IpaA VBSs. We therefore set up to investigate the role of IpaA VBS3 in the formation of vinculin oligomeric complexes induced by IpaA. To this aim, we studied the effects of the IpaA derivatives containing VBS1-2 (A524) or VBS1-3 (A483) on binding to vinculin derivatives (Fig. 2b) using SEC-MALS (Size Exclusion Chromatography-Multi-Angle Light Scattering). Consistent with previous SEC results (Park, Valencia-Gallardo et al. 2011), 3:1 complexes were observed with the first vinculin sub-domain D1 and A483 (Fig. 2c). We then analyzed complexes formed upon incubation of A483 with HV_1-834_ containing the D1-D4 domains, corresponding to full-length human vinculin (HV) devoid of the carboxyterminal F-actin binding domain (Fig. 2b). As shown in Fig. 2d, 1:1 and 2:1 D1D4: A483 heterocomplexes were observed, but not 3:1 complexes (Fig. 2d). Instead and unexpectedly, we detected the formation of a 3:0 D1-D4 homo-trimer (Fig. 2d). Similar 2:1 and 3:0 complexes were observed when A483 was incubated with HV_1-484_ containing only the vinculin D1 and D2 domains (D1D2) (Fig. 2f), indicating that vinculin homo-trimerization only required the vinculin D1 and D2 sub-domains. By contrast, when A524 was incubated with D1D2, 1:1 and 2:1 D1D2:A524 complexes were detected, but not the D1D2 homo-trimer, indicating that IpaA VBS3 was required for vinculin trimerization (Fig. 2e). These results suggest that binding of IpaA VBS1-3 to vinculin triggers conformational changes in vinculin leading to the formation of an IpaA VBS3-dependent vinculin homo-trimer, contrasting with the 3:1 D1-IpaA45-633 heteromeric complex previously observed by SEC analysis using the vinculin D1 subdomain only and suggesting a mechanism different than the proposed scaffolding model by IpaA (Park, Valencia-Gallardo et al. 2011).

**Figure 2.**
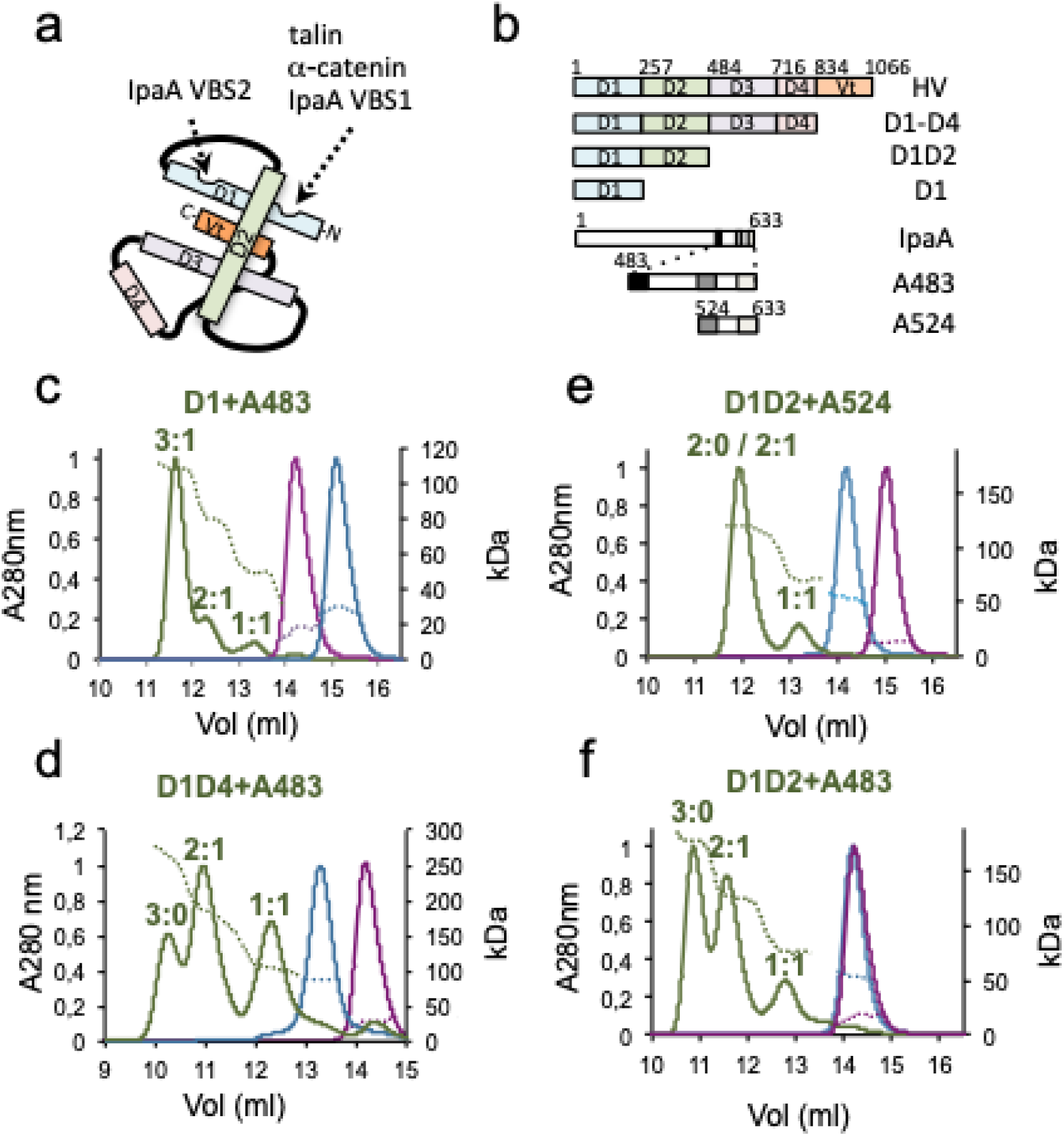
IpaA reveals binding sites in vinculin head subdomains. **a,** Scheme of folded vinculin (HV). The binding sites and corresponding ligands are indicated. **b,** Scheme of HV and IpaA constructs. HV domains and IpaA VBSs are depicted as boxes. The numbers indicate the start residue of each domain. **c-f,** SEC elution profiles of complexes formed between A483 (green) or A524 (purple) and the indicated vinculin derivatives. The indicated complex stoichiometry was inferred from the molecular weight estimated by MALS.

To further investigate initial interactions responsible for vinculin trimer formation, we performed binding assays with HV derivatives immobilized onto a solid phase to restrict conformational changes. These assays indicated that A483 and A524 bound to HV with a similar affinity as estimated by their EC50 (95% confidence interval) of 6.1 (4.2-9.0) and 3.7 (1.7-8.1) nM, respectively (Fig. S2a). Strikingly, a large difference was observed in the binding plateau, indicating that HV presented more binding sites for A483 than for A524 (Fig. S2a). Also, D1D2 presented more binding sites for HV_1-258_ containing the D1 domain only, suggesting the presence of additional sites on the D2 domain (Fig. S2b). Consistently, BN-PAGE showed the formation of 1:1, as well as a 1:2 D1D2:A483 complex, observed with increasing A483 molar ratio (Fig. S2c). In contrast, single 1:1 complexes were observed for D1:A483, D1:A524 or D1D2-A524 (Figs. S2c-h), indicating that IpaA VBS3 was required to reveal additional sites on the D2 domain. Of note, D1D2 higher order complexes observed in the SEC-MALS (Figs. S2c, f-h) were not detected in BN-PAGE, suggesting that Coomassie brilliant blue interfered with the formation of higher order D1D2 complexes. Together, these results suggested that the formation of vinculin trimers triggered by A483 required the IpaA VBS3 dependent exposure of binding sites on D2. These findings were unexpected, since vinculin activating ligands have been described to bind to a single site on the D1 domain of vinculin.

To map interactions of A524 and A483 with D1D2, complexes were cross-linked, subjected to proteolysis and analyzed using Liquid Chromatography coupled to Mass Spectrometry (LC-MS) (Materials and Methods). Intermolecular links were identified from the characterization of cross-linked peptides, and along with identified intramolecular links, used to produce structural models (Suppl. Tables 1-3 and Figs. S3a, 3b; Materials and Methods). The A524:D1 complex showed links consistent with a “canonical” conformer expected from established structures (Izard, Tran Van Nhieu et al. 2006, Tran Van Nhieu and Izard 2007) (Fig. S3c). Similar links were identified for the A524:D1D2 complex, with a majority of links observed with the D1 domain (Fig. S3d). For both complexes, the structure shows interactions between IpaA VBS1 and VBS2 with the D1 first and second bundles, respectively, leading to helical bundle reorganization of D1 associated with vinculin activation (Izard, Evans et al. 2004) (Figs. S3c, d). For the A483:D1D2 complex, MS-based structural modeling reveals two major conformers accounting for the majority of links. In a first “closed” conformer, IpaA VBS1 and VBS2 interact with the D1 bundles in a similar manner as for A524, where the relative positioning of D1 and D2 is globally conserved compared to apo D1D2 or to the A524-D1D2 complex (Fig. 3a and Figs. S3d, e). In this “closed” conformer, IpaA VBS3 interacts with an interface formed by the H5 (residues 128-149) and H8 (residues 222-250) helices in the second bundle of D1, and the H13 (residues 373-397) helix in the second bundle of D2 (Figs. 3a, b). The second “open” conformer, however, shows a major re-orientation of D1 and D2 subdomains with their major axis forming an angle value of ca 82° compared to the 25° observed in the native vinculin structure or the first conformer, with IpaA VBS3 docking sidewise through extensive interaction with the H5 (residues 128-149) and H8 (residues 222-250) helices of D1 (Figs. 3c, d). Since this latter conformer leads to major changes in bundle exposure in D1 and D2 and is only observed for A483, we posit that it is involved in the formation of higher order D1D2 complexes and trimer. To test this, we engineered a structural clamp by substituting residue Q68 in the first D1 bundle and A396 in the second D2 bundle for cysteine residues, expected to prevent the formation of the open conformer upon disulfide bridge formation (Fig. 3e). Consistent with the SEC-MALS analysis, higher order D1D2 homo-complexes devoid of A483 and D1D2:A524 hetero-complexes were visualized by native PAGE migrated in the absence of Coomassie brilliant blue (Fig. 3f and Figs. S4c, d). The cysteine clamp Q68C A396C (CC) in D1D2 did not prevent the exposure of additional sites on D2 or 1:1 complex formation induced by A524 or A483. However, CC prevented the formation of higher order complexes (Figs. 3e, f and Fig. S4). Accordingly, we coined “supra-activation” the mode of vinculin activation induced by A483 involving major conformational changes in the vinculin head, to distinguish it from the canonical activation associated with the dissociation of vinculin head-tail domains.

**Figure 3.**
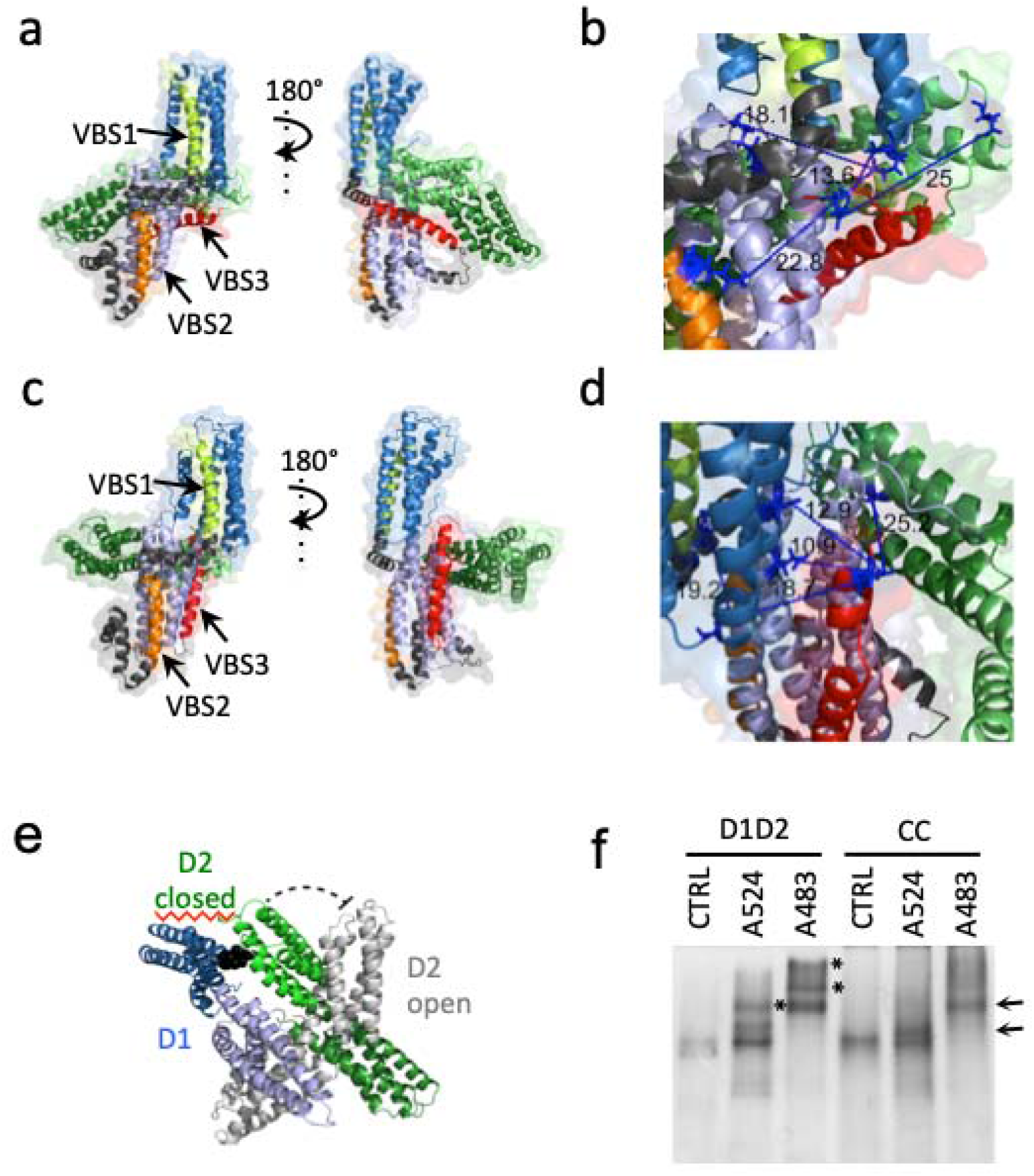
Characterization of IpaA contact sites on vinculin. **a-d**) Structural models of D1D2-A483. **a, b,** “closed” conformer; **c, d,** “open” conformer. **b, d,** higher magnification of the IpaA VBS3-D1D2 interaction in (**a**) and (**c**) showing the identified cross-linked distance between residues in Å. IpaA VBS1-3 were docked on the surface of Vinculin D1D2 and verified using MS cross-link constraints. TX-MS protocol (Hauri, Khakzad et al. 2019) in combination with MS constraints was used to unify and adjust the final model, which justifies over 100 cross-links. **e,** Structural model of cysteine-clamped vinculin. Blue: D1 domain. Green: D2 domain in the closed conforme. Grey: D2 domain in the open conformer. Black: C68-C396 cysteine clamp preventing the switch from closed to open conformers. **f,** native gel analysis of vinculin D1D2 and IpaA derivatives. D1D2 or cysteine-clamped D1D2 (CC) were incubated with the indicated IpaA derivatives and analyzed by native PAGE followed by Coomassie staining. Arrows: 1:1 complexes. *: higher order complexes. Note the absence of higher order complexes for cysteine-clamped D1D2.

Because bacterial pathogens often exploit host cell endogenous processes, we next tested the effects of vinculin supra-activation on FA formation by introducing the cysteine clamp in full length vinculin fused to mCherry (CC-HV) and analyzed its effects following transfection in MEF vinculin-null cells. As shown in Fig. 4a, CC-HV led to the formation of larger and more numerous talin-containing FAs than mock-transfected vinculin null cells, consistent with residual vinculin activation (Figs. 4a, b). In contrast, CC-HV-expressing cells formed significantly fewer and smaller FAs than cells transfected with wild-type vinculin when analyzed for talin or vinculin (Figs. 4a-e). A more detailed analysis indicated that CC-HV formed adhesion microclusters remarkably conserved in width with an average of 0.96 ± 0.27 (SD) μm, connected to actin fibers (Figs. 4f, h, i). This was in sharp contrast with FAs formed by wild-type vinculin showing an average width of 1.3 ± 0.27 (SD) μm and reaching up to several microns (Figs. 4g, h, i). These results suggest that vinculin supra-activation and oligomerization, impaired in CC, is involved in the merging of adhesion microclusters during FA maturation. These findings also raised the issue about the role of IpaA-mediated vinculin supra-activation in the context of endogenous adhesion processes. To address this, we compared the effects of A524 and A483 expression on FAs. As shown in Figs. 4j-l and S5a, b, cells transfected with GFP-A524 formed more numerous and larger peripheral FAs as well as actin-rich ruffles compared to control cells. Strikingly, GFP-A483 transfected cells formed even larger and more numerous FAs and significantly less actin ruffles than GFP-A524 transfected cells (Figs. 4j-l). Furthermore, GFP-A483-induced FAs were extremely stable, with a median duration of at least 84 min, while GFP-A524 and control cells showed FAs with a comparable median duration of less than 25 min (Figs. S5c, d; Suppl. movie 1). This increased FA stability in GFP-A483 transfectants was predominantly due to decreased rates of FA disassembly, with a 2-fold decrease in median instant rates relative to control cells (Figs. S5e, f; Suppl. movie 1).

**Figure 4.**
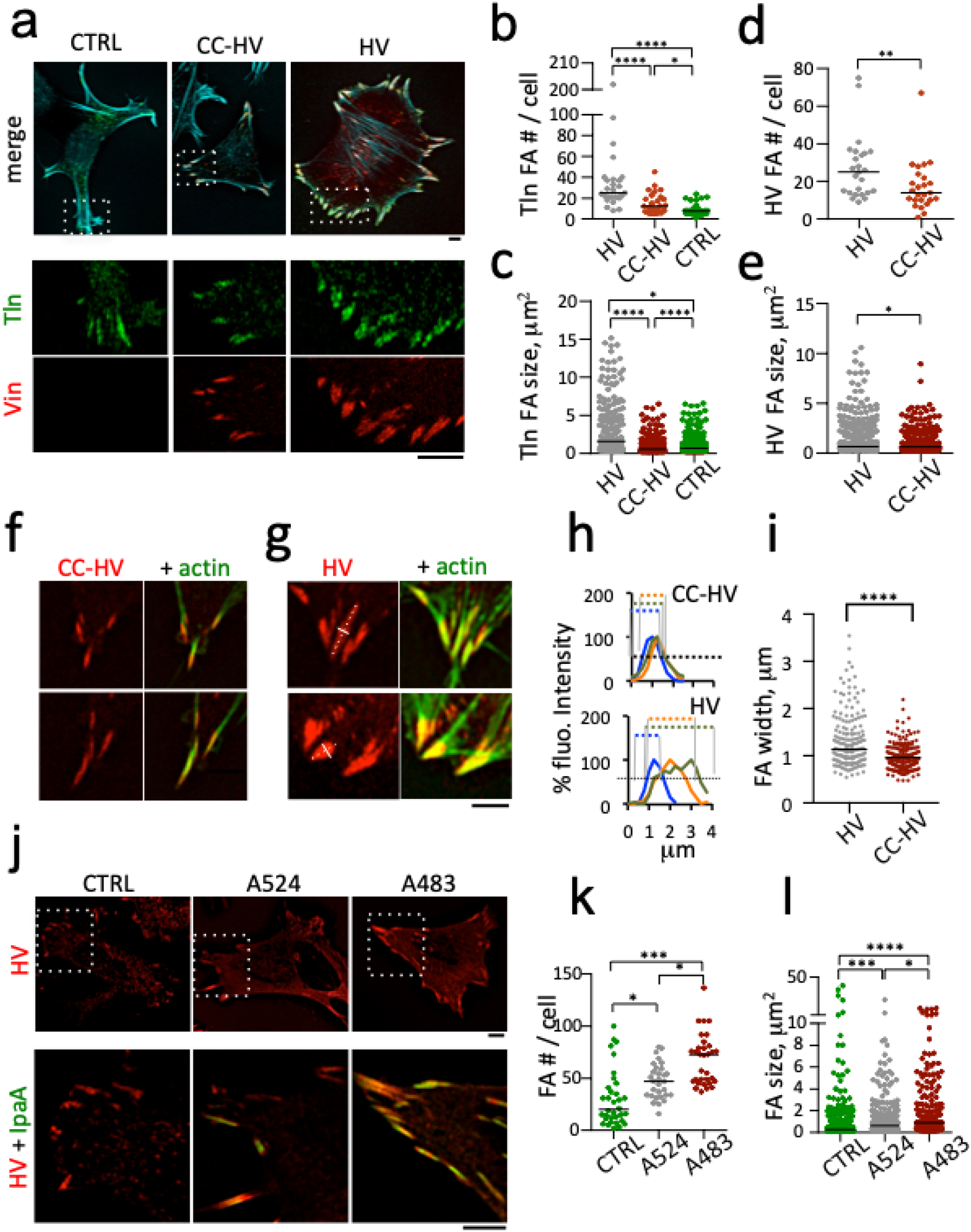
vinculin supra-activation promotes the merging of adhesions clusters. Cells were transfected, fixed and processed for fluorescence analysis of FAs. **a, f, g, j,** representative fluorescence micrographs; red: Vinculin-mCherry; green: GFP-talin (**a**) or F-actin (**f, g**); cyan: F-actin (**a**). Scale bar = 5 μm. **a-i,** MEF vinculin null cells; transfection with GFP-talin (CTRL); talin co-transfection with full-length vinculin-mCherry (HV) or HV Q68C A396C-mCherry (CC-HV). **j-l,** C2.7 cells; transfection with HV (CTRL); HV co-transfection with GFP-A524 or GFP-A483. **b-e, k, l**: the FA number per cell and size were determined using a semi-automatic detection program (Materials and Methods). Bar: median size. FAs analyzed for: **b, c,** GFP-talin; **d, e, h, i, k, l,** HV or CC-HV. **b-e,** CTRL: n=28, N=3; HV: n = 25, N = 3; CC-HV: n = 25, N = 3. **h,** representative plot profiles from linescans of FA width as depicted in **(g)** by the solid white line orthogonal to the main FA axis (dashed white line); **g, i,** FA width determined as the full width half maximum by linear interpolation from plot profiles in (**h**); HV: 181 FAs, 6 cells, N = 2. CC-HV: 101 FAs, 14 cells, N = 3. Mann-Whitney test with Bonferroni multiple comparison correction. *: p < 0.05; **: p < 0.01; ***: p < 0.005; ****: p < 0.001. **j-l,** n > 30 cells, N = 3. Dunn’s multiple comparisons test. *: p < 0.05; ***: p < 0.005.

Strikingly, GFP-A483-induced FAs resisted the action of the Rho-kinase inhibitor Y27632 relaxing actin-myosin, with a five- and four-times slower median rate of FA disassembly relative to control cells and GFP-A524 transfectants, respectively (Figs. 5a-d; Suppl. movie 2). Large FAs were even observed to form in GFP-A483 transfectants following addition of the inhibitor (Figs. 5a, c, d), a process that was not observed for other samples, including cells transfected with GFP fused to the vinculin D1 domain (vD1) reported to delay talin refolding following stretching (del Rio, Perez-Jimenez et al. 2009, Margadant, Chew et al. 2011, Carisey, Tsang et al. 2013) (Figs. 5a, b, d; Suppl. movie 2). GFP-A483 also delayed the Y27632-induced removal of the late adhesion marker VASP (Figs. 5e-g; Suppl. movie 3). Together, our findings suggest that vinculin supra-activation occurs during FA maturation but is triggered by A483 in the absence of mechanotransduction. To further investigate this, we performed cell replating experiments. GFP-A483 induced higher yields of adherent cells when replating was performed with short kinetics, with a 5-fold increase over control cells and GFP-A524 transfectants, respectively, for 10 min replating (Fig. 6a). By contrast, little difference in adhesion yield was detected between samples at 15 min suggesting that IpaA predominantly affected the early dynamics of cell adhesion (Fig. 6a). To extend these findings, we measured cell adhesion strength using controlled shear stress in a microfluidic chamber and 1025 Lu melanoma cells (Smalley, Lioni et al. 2008). We first controlled the efficacy of anti-vinculin siRNA treatment in these cells (Figs. 6b, c). Consistent with replating experiments, when cells were allowed to adhere to fibronectin-coated surfaces for more than 25 min, little difference in resistance to shear stress could be detected between GFP-A483 and GFP transfected cells samples (Figs. 6d, e). In contrast, similar to cells depleted for by siRNA treatment, cells transfected with the clamped vinculin version showed a decreased ability to adhere in comparison to wild-type vinculin-transfected cells (Fig. 6e). However, when shear stress was applied after less than 20 min following cell incubation, GFP-A483-transfected cells showed significantly higher resistance to shear stress up to 22.2 dynes.cm^−2^ than GFP-A524- or GFP-transfected cells, with 1.7 ± 0.2 -and 0.9 ± 0.14-fold enrichment ± SD of adherent cells for GFP-A483 and GFP-A524-transfected cells versus control GFP-transfected cells, respectively (Fig. 6f; Suppl. movie 4). These results are in full agreement with effects observed on adhesion structures and suggest that A483-mediated vinculin supra-activation accelerates endogenous processes occurring during mechanotransduction to promote strong adhesion.

**Figure 5.**
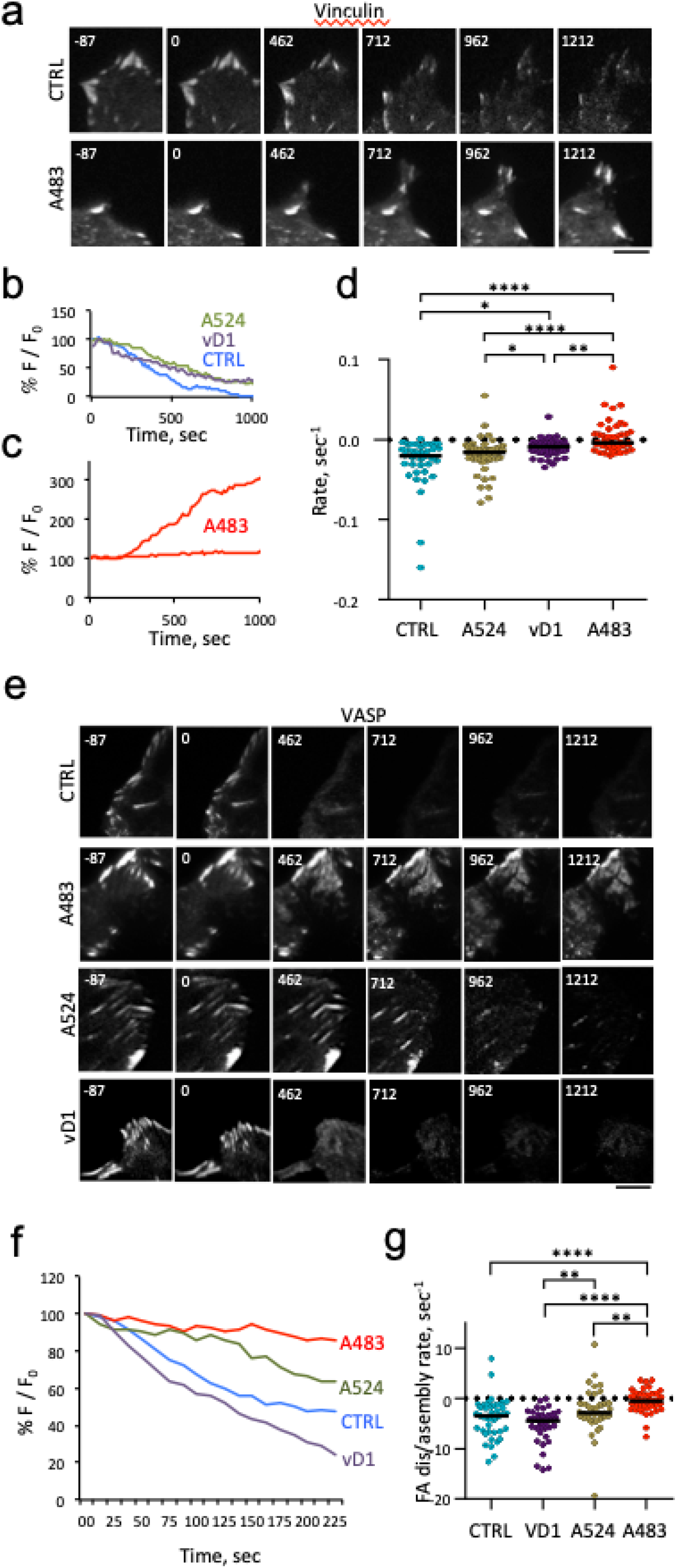
IpaA-mediated vinculin supra-activation stimulates cell adhesion independent of mechanotransduction. TIRF microscopy of C2.7 cells transfected with mCherry-vinculin **(a-d)** or mCherry-VASP **(e-g)** alone (CTRL) or co-transfected with GFP-IpaA VBS1-2 (A524), GFP-vD1 (vD1), or GFP-IpaA VBS1-3 (A483). Adhesion kinetic parameters were determined from time-lapse acquisitions following cell treatment with 100 μM Y-27632. **a, e,** representative time series acquisitions. Numbers indicated the elapsed time in seconds, with the inhibitor added at t = 0. Scale bar = 5 μm. **b, c, f,** % F/F_0_: average fluorescence intensity of adhesions expressed as a percent of initial fluorescence. Representative traces corresponding to single adhesions for the indicated samples. **d, g,** initial rates of adhesion assembly / disassembly inferred from linear fits. Number of adhesions analyzed: **d,** N = 5. CTRL: 84; vD1: 75; A524: 140; A483: 97. **f,** N = 4. CTRL: 42; A483: 43; A524: 40; vD1: 40. Dunn’s multiple comparisons test. *: p < 0.05; ****: p < 0.001.

**Figure 6.**
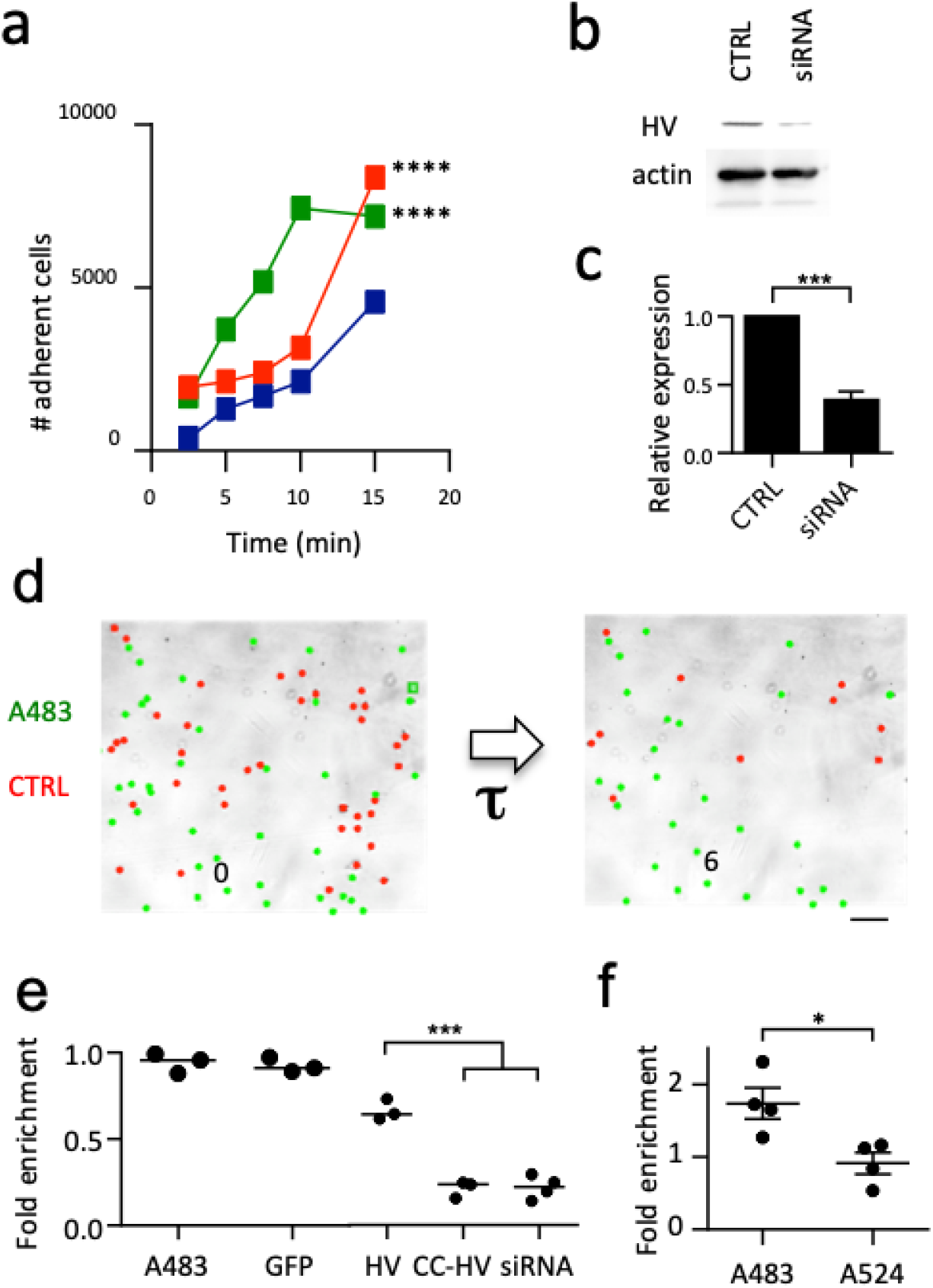
IpaA increases the kinetics of cell adhesion and strength. **a,** 1205Lu melanoma cells were transfected with GFP alone (blue), GFP-IpaA VBS1-2 (A524, red or GFP-IpaA VBS1-3 (A483, green), lifted up by trypsinization and plated for the indicated time on Fn-coated coverslips. Samples were washed, fixed and adherent cells were scored microscopically. The total number of adherent cells scored is indicated. GFP: 3223 cells, N = 4; A524: n=7418, N = 4; A483: n = 5668, N = 4. Chi square corrected with Bonferroni multiple comparison correction. ****: p < 0.001. **b, c,** 1205Lu melanoma cells were mock-transfected (CTRL) or treated with anti-vinculin siRNA (siRNA, Materials and Methods). **b,** anti-vinculin Western blot analysis. **c,** average HV band intensity normalized to that of control cells. Unpaired t test. ***: p = 0.005. **d-f,** transfected cells were with calcein (Materials and Methods) and mixed with the same ratio of control cells. Cells were perfused in a microfluidic chamber and allowed to adhere prior to shear stress application for: **d, e**: 30-60 min; **f,** 20 min. **d,** representative fields. The number indicates the elapsed time (seconds) after shear stress application. **e, f,** scatter plot of the ratio of adherent cells with respect to non-transfected cells, **e,** A483 (N = 3, n = 557); GFP (N = 3, n = 490); HV: vinculin mCherry (N = 3, n = 481); CC-HV: vinculin Q68C A396C-mCherry (N = 3, n = 259); siRNA: cells treated with anti-vinculin siRNA (N = 3, n = 395). **f,** A483 (N = 4, n = 610) or A524 (N = 4, n = 433) transfected cells vs control cells (1594 cells, N = 4). Unpaired t test. *: p = 0.0229.

Vinculin is paradoxically described as a prognostic marker favoring the migration of cancer cells or as a tumor suppressor stimulating cell anchorage (Goldmann, Auernheimer et al. 2013, Labernadie, Kato et al. 2017, Hamidi and Ivaska 2018). These contradictory findings reflect its complex and poorly understood regulation, as well as different roles in 2D or 3D systems (Mierke, Kollmannsberger et al. 2010, Gulvady, Dubois et al. 2018). Also, an increase of the total pool of vinculin may not correlate with increased vinculin activation. We took advantage of the unique property of A483 to study the effects of vinculin supra-activation on the motility and invasion of melanoma cells. In time-lapse microscopy experiments in 2D-chambers, GFP-A524 inhibited melanocyte motility compared to control cells, with a rate of Root Median Square Displacement (rMSD) of 3.16 and 15.6 μm.min^−1^, respectively (Figs. 7a, b). An even stronger inhibition was observed for GFP-A483-transfected cells (rMSD = 2.3 μm.min^−1^) (Figs. 7a, b). Transmigration of melanocytes in 3D-matrigels was similarly inhibited by A524 and A483 (Fig. 7c), demonstrating the potential of A483 to counter the invasiveness of tumor cells. Such efficient inhibition of cell migration by IpaA is expected from its ability to trigger adhesion formation in the absence of actomyosin contraction.

**Figure 7.**
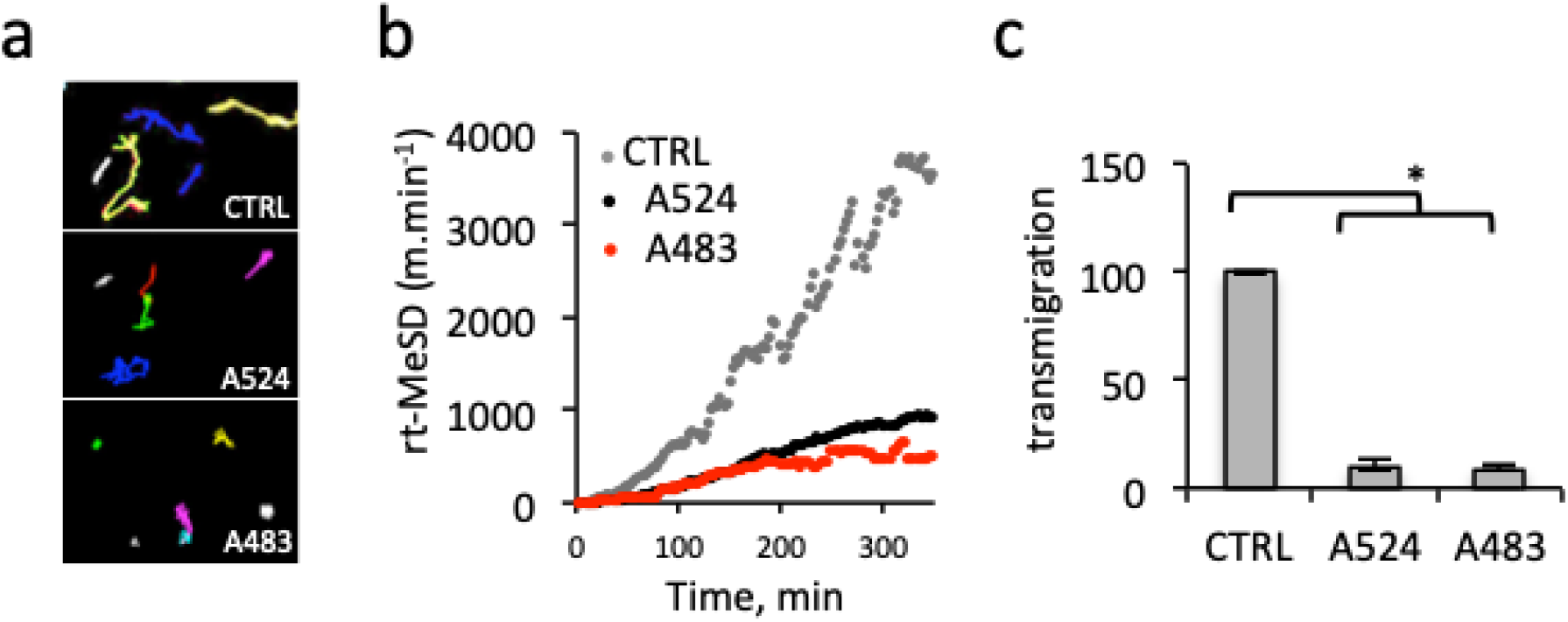
A483 inhibits tumor cell invasion. 1205Lu melanoma cells were transfected with GFP alone (blue), GFP-IpaA VBS1-2 (A524, red) or GFP-IpaA VBS1-3 (A483, green), lifted up by trypsinization and plated for the indicated time on Fn-coated coverslips. Samples were transfected with the indicated constructs, and analyzed by time-lapse videomicroscopy; **a,** representative single cell migration 20-hour tracks for indicated samples. **b,** Root of Median Square of displacement over time for control- (61 cells, N = 3), A524- (61 cells, N = 3) and A483 transfectants (64 cells, N = 3). ***: p = 0.0007. The slopes were analyzed using a covariance test and found to be statistically different (ANCOVA, p < 2×10^−16^). **c,** 5 × 10^4^ cells were seeded in matrigel chambers. The percent of transmigrated cells is indicated. (N = 3). Kruskal-Wallis test with Dunn’s multiple comparisons test. *: p < 0.05.

## Discussion

Through the joint action of its VBSs, IpaA induces major conformational changes of the vinculin head, unveiling binding sites on the D2 subdomain. Vinculin head subdomain exposure was observed in molecular dynamics stretching simulations (Kluger, Braun et al. 2020), suggesting its occurrence during mechanotransduction. Interestingly, IpaA VBS3 that binds to D2 shares homology with the talin VBS corresponding to helix 46, suggesting that this talin VBS combined with actomyosin stretching forces could promote vinculin supra-activation. In the case of IpaA, the lack of requirement for mechanotransduction may be linked to the extreme affinity of IpaAVBS1, 2 for vinculin D1 (Tran Van Nhieu and Izard 2007), insuring the stable positioning of the IpaAVBS2-3 linker region onto D1D2 and enabling IpaA VBS3 interaction with D2. Our in vitro experiments indicate that unlike vinculin canonical activation, IpaA supra-activation leads to vinculin oligomerization via Vh-Vh interactions. Consistent with a role for vinculin supra-activation, cysteine-clamped vinculin fails to form oligomers and forms smaller adhesions of restricted width (Fig. 4h). These observations suggest that vinculin canonical activation induces adhesion microclusters, but that vinculin supra-activation is required for the merging of these microclusters. Our in results indicate that merging of microclusters is associated with the formation of high order vinculin oligomers. While Vt-Vt interaction is known to induce dimerization, its combination with D1D2-D1D2 interactions could provide the basis for the formation of high order vinculin complexes (Fig. 8). Along these lines, the initially observed “parachute” complexes were proposed to correspond to up to six vinculin molecules forming through Vh-Vh interactions and connected via their tails, which, in view of our findings, could correspond to a pair of vinculin trimers (Molony and Burridge 1985). Such high order vinculin structures could participate in the scaffolding of focal adhesion components and clustering of adhesion structures. Clustering of adhesions at different scales is believed to play a major role in adhesive processes through the regulation of functional units (Mege and Ishiyama 2017). The basis for integrin clustering in adhesions is not fully understood and may be induced by ligand-binding or integrin homo-oligomerization via integrin trans-membrane domains (Karimi, O’Connor et al. 2018). At the nanoscale, integrin nanoclusters were proposed to correspond to elementary units merging to form nascent adhesions (Changede and Sheetz 2017). At a larger scale, vinculin supra-activation provides a basis for the merging of adhesion microclusters into larger adhesion structures (Fig. 8).

**Fig. 8.**
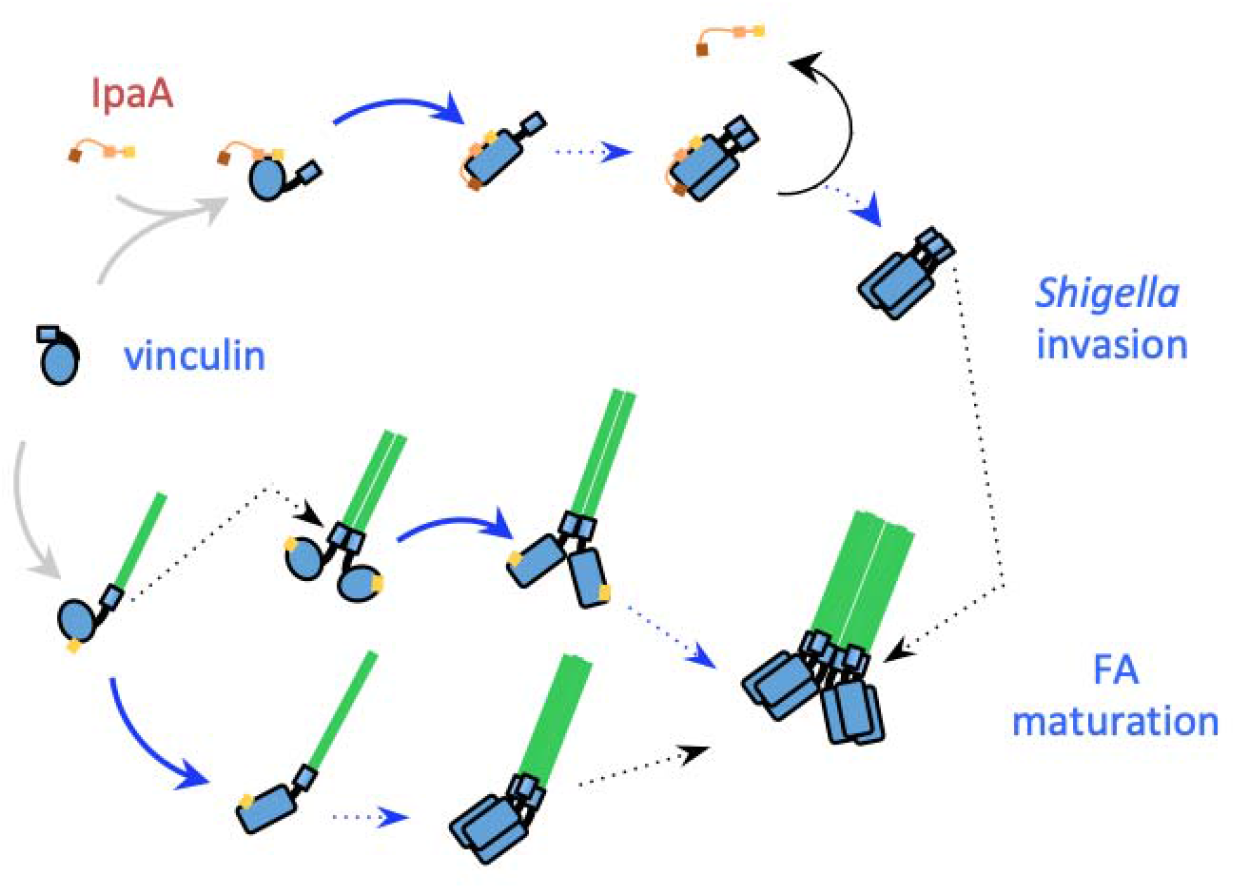
Scheme of IpaA-mediated vinculin supra-activation and oligomerization during *Shigella* invasion and FA maturation. Vinculin activation leading to disruption of Vh-Vt interaction (grey arrows) occurs following binding of IpaA VBS1, 2 to D1 or binding of an endogenous VBS to D1. Supra-activation (blue arrows) is triggered by the concerted action of the three IpaA VBS1-3 independent of mechanotransduction (top), and may occur during mechanotransduction (bottom). Supra-activation leads to major conformational changes in Vh and vinculin oligomerization through Vh-Vh interaction (blue dotted arrows) resulting in the formation of high order vinculin complexes when combined with Vt-Vt dimerization (black dotted arrows). Green, actin fibers; blue circle: Vh; blue rectangle: Vt; yellow: activating VBS; orange: IpaA VBS2; red: IpaA VBS3. 1): mechanotransduction-independent. 2): mechanotransduction-dependent.

IpaA may act as a catalyst, released and recycled from vinculin complexes during vinculin homo-trimer formation, potentially accounting for the potency of its effects. *Shigella* invading through a triggering mode relies on a discrete number of host cell contacts for which cytoskeletal tethering is likely critical for invasion (Valencia-Gallardo, Carayol et al. 2015). As opposed to physiological substrates, bacteria cannot sustain the range of counter-forces associated with integrin-mediated adhesion to the substrate. The *Shigella* type III effector IpaA provides an elegant solution to this problem by promoting strong adhesion without requirement for mechanotransduction. Understanding how these major vinculin conformational changes regulate the composition and properties of cell adhesions will bring important insights into cell adhesion processes and will be the focus of future investigations.

## ACKNOWLEDGEMENTS

The authors thank Gauthier Mercante for technical help, Philippe Mailly for help with image analysis and René-Marc Mège for insightful discussions and reading of the manuscript. This work was supported by grants from Inserm, CNRS and Collège de France to the CIRB, as well as grant from the PSL Idex project “Shigaforce”. DIAS and BCC are recipients of a PhD fellowship from a CONACYT scholarship. CV-G and DIAS also received support from the Memolife Labex. HK and LM were supported by Swiss National Science Foundation (grant no. SNF 200021 160188) and by SNF P2ZHP3_191289 and the Knut And Alice Wallenberg Foundation (grant no. KAW 2016.0023), respectively.

## AUTHOR CONTRIBUTIONS

CV-G and DIAS conceived and performed most of the experimental works, data analysis and wrote the manuscript. BCC and BM analyzed TIRF experiments. CB-N performed the SEC-MALS analysis. CB and NQD performed and analyzed experiments with melanocytes with the help of DJ and AM. BCC, AA and JF provided technical help for cell adhesion and microfluidics experiments. CM and JCR designed and performed the LC-MS analysis. DBL analyzed the cross-linked mass spectrometry data. HK and LM generated structural models. GTVN designed the project and wrote the manuscript.

## Materials and Methods

### Plasmids and constructs

Human vinculin constructs were generated by polymerase chain reaction using the forward primer 5’ GCGCATATGCCAGTGTTTCATACG-3’ and reverse primers 5’-CGTCGACTCACCAGGCATCTTCATCGGC-3’ for D1 (residues 1-258) or 5’-CGTCGACTCAGTGTACAGCTGCTTTG-3’ for D2 (residues 1-492) using a plasmid containing full-length octahistidine-tagged human vinculin (residues 1–1,066), as template (Bakolitsa, Cohen et al. 2004), and cloned into the NdeI-SalI sites of pet15b (Novagen) to obtain pET15b-D1 and pET15b-D1D2, respectively. The Q68C and A396C cysteine substitution for the cysteine clamp were introduced into pet15b-D1D2 by site-directed mutagenesis using the 5’- GAGACTGTTCAAACCACTGAGGATTGCATTTTGAAG-3’ and 5’- ATCGATGCTGCTCAGAACTGGCTTTGCGATCCAAAT-3’ primers, respectively. The pGFP-vD1 plasmid was generated by polymerase chain reaction using the forward primer 5’-ACCCGGGATCCCGCC-3’ and reverse primer 5’-ACCCGGGACCAGGCA-3’, and cloned into peGFP. The pmCherry-human vinculin (HV) and pmCherry-VASP plasmids were from Addgene. Stealth siRNA anti-human vinculin was from Invitrogen (reference number 1299001). The cysteine clamp was introduced in pmCherry N1-HV by exchanging the NheI-PspXI fragment with the corresponding XbaI-PspXI fragment of pET15b-D1D2 -Q68C A396C.

The IpaA constructs GFP-A524 and GFP-A483 were generated by polymerase chain reaction (PCR) and cloning into pcDNA3.1 NT-GFP Topo TA (Invitrogen) using the 5’-TCAAAGGACATTACAAAATCC-3’ and 5’-GCGATATCATGGCCAGCAAAGG-3’ forward primers, respectively, and the 5’- GCGCGGCCGCTTAATCCTTATTGATATTC-3’ reverse primer. The GST-A483A construct was generated by PCR using 5’-GGCGAATTCCCGGAGACACATATTTAACACG-3’ forward and 5’- GCCGTCGACTTAATCCTTATTGATATTCT-3’ reverse primers and cloning into the *EcoRI-SalI* ofpGEX-4T-2 (GE Lifesciences). pGST-A524 was previously described (Ramarao, Le Clainche et al. 2007). The pGFP-vD1 plasmid was generated by polymerase chain reaction using the forward primer 5’-ACCCGGGATCCCGCC- 3’ and reverse primer 5’-ACCCGGGACCAGGCA-3’, and cloned into peGFP. The pmCherry-human vinculin (HV) and pmCherry-VASP plasmids were from Addgene. Stealth siRNA anti-human vinculin was from Invitrogen (reference number 1299001). All constructs were verified by DNA sequencing.

### Cell lines and bacterial strains

HeLa cells (ATCC CCL-2) were incubated in RPMI (Roswell Park Memorial Institute) medium containing 5% FCS (fetal calf serum, Gibco^®^) in an incubator with 5% CO_2_. C2.7 myoblasts (Mitrossilis, Fouchard et al. 2009) and MEF vinculin null cells (Humphries, Wang et al. 2007) were routinely grown in DMEM 1 g / L glucose containing 10 % FCS in a 37°C incubator containing 10 % CO_2_. 1205Lu melanoma cells (Smalley, Lioni et al. 2008) were grown in RPMI + Glutamax medium (RPMI1640) supplemented with 10 % fetal calf serum (FCS) and non-essential aminoacids in a 37°C incubator with 5 % CO_2_. For transfection experiments, cells were seeded at 2.5 × 10^4^ cells in 25 mm-diameter coverslips. Cells were transfected with 3 μg of pGFP-A524 or pGFP-A483 plasmids with 6 μls JetPEI transfection reagent (Polyplus) for 16 hours following the manufacturer’s recommendations. C2.7 mice myoblasts cells were fixed in PBS containing 3.7% paraformaldehyde for 20 min at 21°C and permeabilized with 0.1% Triton X-100 for 4 min at 21°C. 1205Lu melanoma cells were processed for adhesion under shear stress experiments in microfluidic chambers.

The wild type *Shigella flexneri*, isogenic mutants, and complemented *ipaA* mutant strains, as well as wild type *Shigella* expressing the AfaE adhesin were previously described (Izard, Tran Van Nhieu et al. 2006). Bacterial strains were cultured in trypticase soy broth (TCS) medium at 37°C. When specified, antibiotics were added at the following concentrations: carbenicillin 100 μg/ml, kanamycin 20 μg/ml.

### Cell challenge with *Shigella* strains

HeLa cells seeded at 4 × 10^5^ cells in coverslip-containing 34 mm-diameter wells the day before the experiment. After 16 hours, cells were challenged with *Shigella* strains coated with poly-L-lysine, as follows. Bacteria grown to an OD_600 nm_ of 0.6 - 0.8 were washed three-times by successive centrifugation at 13 Kg for 30 sec and resuspension in EM buffer (120 mM NaCl, 7 mM KCl, 1.8 mM CaCl_2_, 0.8 mM MgCl_2_, 5 mM glucose, and 25 mM HEPES, pH = 7.3). Samples were resuspended in EM buffer containing 50 μg/ml poly-L-lysine and incubated for 15 min at 21°C, washed three times in EM buffer and resuspended in the same buffer at a final OD of OD_600 nm_ = 0.2. Cell samples were washed three times in EM buffer and challenged with 1 ml of the bacterial suspension and incubated at 37°C. Samples were fixed with PBS containing 3.7% PFA after 30 min incubation. Samples were processed for immunofluorescence microscopy.

### Immunofluorescence staining

Cells were processed for immunofluorescence staining using the Vin11.5 anti-vinculin monoclonal antibody (ref. V4505, Sigma-Aldrich) and anti-mouse IgG antibody coupled to Alexa 546 (Jackson Research) and Phalloidin-Alexa 633 (Invitrogen), as described previously (Tran Van Nhieu and Izard 2007). Bacteria were labeled using anti-LPS rabbit polyclonal antibody followed by anti-rabbit IgG antibody coupled to Alexa 525 as described (Izard, Tran Van Nhieu et al. 2006). Samples were analyzed using an Eclipse Ti inverted microscope (Nikon) equipped with a 60 x objective, a CSU-X1 spinning disk confocal head (Yokogawa), and a Coolsnap HQ2 camera (Roper Scientific Instruments), controlled by the Metamorph 7.7 software.

### Protein purification

BL21 (DE3) chemically competent *E. coli* (Life Technologies) was transformed with the expression constructs. D1 and D1D2 were purified essentially as described (Papagrigoriou, Gingras et al. 2004, Park, Valencia-Gallardo et al. 2011). For the IpaA derivatives, bacteria grown until OD_600nm_ = 1.0 were induced with 0.5 mM IPTG and incubated for another 3 hrs. Bacteria were pelleted and washed in binding buffer 25 mM Tris PH 7.4, 100 mM NaCl and 1 mM beta-mercaptoethanol, containing Complete™ protease inhibitor. Bacterial pellets were resuspended in 1/50th of the original culture volume and lyzed using a cell disruptor (One shot model, Constant System Inc.). Proteins were purified by affinity chromatography using a GSTrap HP affinity column (GE Healthcare) and size exclusion chromatography (HiLoad S200, Ge Healthcare). Samples were stored aliquoted at −80°C at concentrations ranging from 1 to 10 mg/ml.

### Protein complex formation analysis

Proteins were incubated at a concentration of 30 μM in binding buffer for 60 min at 4°C. Samples were analyzed by SEC-MALS (Wyatt Technology Europe) using a 24 ml Superdex 200 Increase 10/300 GL filtration column and a MiniDAWN TREOS equipped with a quasi-elastic light scattering module and a refractometer Optilab T-rEX (Wyatt Technology). Data were analyzed using the ASTRA 6.1.7.17 software (Wyatt Technology Europe). Protein complex formation was visualized by PAGE under non-denaturing conditions using à 7.5% polycrylamide gel, followed by Coomassie blue staining.

### Solid-phase binding assay

96–well Maxisorp (Nunc) ELISA plates were coated with 30 nM of full-length vinculin, vinculin constructs or IpaA proteins at the indicated concentrations in binding buffer (25 mM Tris PH 7.4, 100 mM NaCl and 1 mM β-mercaptoethanol). Samples were blocked with PBS-BSA 2%, washed and incubated with IpaA or vinculin proteins in binding buffer containing 0.2% BSA at room temperature for one hour. After incubation, the plates were washed and incubated with an anti-IpaA (dilution 1/2000^e^) polyclonal primary antibody^3^ or anti-vinculin (dilution 1/2000^e^) Vin11Vin.5 monoclonal antibody (Sigma-Aldrich) in binding buffer containing 0.2% BSA for one hour at room temperature. Plates were washed and incubated with an HRP-coupled secondary anti-rabbit or anti-mouse IgG antibody (1/32000^e^) (Jackson ImmunoResearch) for one hour. The reaction was revealed by adding 100 μl of tetramethylbenzidine (Sigma-Aldrich) for 15 min, stopped by adding 50 μl of 0.66N H_2_SO_4_ and the absorbance was read at 450 nm (Dynatech MR400).

### BN-PAGE (Blue Native – Polyacrylamide Gel Electrophoresis) protein native gel analysis and complex cross-linking

25 μM of vinculin constructs were incubated with different molar ratios of IpaA proteins in a 1X BN-PAGE buffer (250 mM ∊-aminicaprionic acid and 25mM Bis-Tris PH 7,0) at 4°C for one hour. The protein mixtures were separated in a one-dimension native BN-PAGE electrophoresis as described (Eubel and Millar 2009). For vinculin-IpaA protein ratio assay, vinculin-IpaA bands containing the complexes separated by BN-PAGE were cut, sliced and boiled in 2 x Laemmli SDS buffer followed by SDS-PAGE. The second dimension SDS-PAGE gels were stained (colloidal Coomassie staining) and the density of the bands was determined using Image J. The normalized vinculin:IpaA ratio of the complexes was compared using a non-parametric Kruskal-Wallis rank sum test (R statistical software).

For crosslinking vinculin-IpaA complex, bands containing the complexes were cut, sliced and electroeluted in native conditions (15 mM Bis-Tris pH 7.0 and 50 mM Tricine) inside a closed dialysis membrane (SpectraPor). The soluble complexes were recovered and their buffer exchanged twice into an amine-free cross-link buffer in 25 mM HEPES pH 7.0 containing 100 mM NaCl using 10MWCO ZEBA desalting columns (Thermo Scientific). The fractions containing the complexes were incubated for 1 hr at 4°C with 10 mM N-hydroxysulfosuccinimide and 5 mM EDC (Sigma-Aldrich) following the manufacturer’s recommendations. The cross-linking reaction was stopped by adding 50 mM Tris pH 7.4 and incubating for 20 minutes. Samples were denaturated in 2x SDS Laemmli buffer for 5 min at 95°C and complexes were eluted from gel slices following SDS-PAGE.

### Liquid Chromatography Mass spectrometry (LC-MS)

Complexes obtained after the cross-linking step were loaded onto a 4-20% polyacrylamide gradient gels and Coomassie stained. The bands containing the complexes were cut and submitted to tryptic digestion (Shevchenko, Tomas et al. 2006). The experiments were performed in duplicates for the 3 complexes D1:A524, D1D2:A524 and D1D2:A483. Peptides obtained after tryptic digestion were analyzed on a Q Exactive Plus instrument (Thermo Fisher Scientific, Bremen) coupled with an EASY nLC 1 000 chromatography system (Thermo Fisher Scientific, Bremen). Sample was loaded on an in-house packed 50 cm nano-HPLC column (75 μm inner diameter) with C18 resin (1.9 μm particles, 100 Å pore size, Reprosil-Pur Basic C18-HD resin, Dr. Maisch GmbH, Ammerbuch-Entringen, Germany) and equilibrated in 98 % solvent A (H2O, 0.1 % FA) and 2 % solvent B (ACN, 0.1 % FA). A 120 minute-gradient of solvent B at 250 nL.min-1 flow rate was applied to separate peptides. The instrument method for the Q Exactive Plus was set up in DDA mode (Data Dependent Acquisition). After a survey scan in the Orbitrap (resolution 70 000), the 10 most intense precursor ions were selected for HCD fragmentation with a normalized collision energy set up to 28. Charge state screening was enabled, and precursors with unknown charge state or a charge state of 1 and >7 were excluded. Dynamic exclusion was enabled for 35 or 45 seconds respectively.

### Data analysis

The identification of cross-linked peptides from LC-MS data was performed using SIM-XL v. 1.3 (Lima, de Lima et al. 2015), with the following search parameters: EDC as cross-linker, a tolerance of 20 ppm for precursor and fragment ions, trypsin fully specific digestion with up to three missed cleavages. Carbamidomethylation of cysteines was considered as a fixed modification. All initial identification of cross-linked peptides required a primary score of SIM-XL greater than 2.5 for inter-links and 2.0 for intra-links or loop-links. As single incorrect cross-link identification might lead to a different model, a manual post-validation of the search engine results at the MS2 level was thus performed. A 2D-map showing the protein-protein interaction was generated as an output (Figs. 2a,b). Only peptides present in the 2 replicates are gathered in Supplementary Tables 1–3 and were used for the modeling.

### Modeling

We used the distance constraints obtained from cross-linking MS data (Suppl. Tables 1–3) to guide the protein structure modeling using the TX-MS protocol as described by Hauri, Khakzad et al. (Hauri, Khakzad et al. 2019). In short, TX-MS uses the Rosetta comparative modeling protocol (RosettaCM) (Song, DiMaio et al. 2013), and the flexible backbone docking protocol (RosettaDock) (Gray 2006) to generate models and evaluate how well each model explains the MS constraints using a novel scoring function. Here, a total of 100,000 models was generated, of which the highest-scoring model is displayed in (Fig. 3c), supported by a total of 100 inter and intra-molecular cross-links.

### TIRF (Total Internal Reflection Microscopy) analysis

C2.7 cells were transfected with pmCherry-HV or pmCherry-VASP and the indicated plasmids as described above. Samples were mounted onto a TIRF microscopy chamber on a stage of an Eclipse Ti inverted microscope (Nikon) equipped with an Apo TIRF 100 x N.A. 1.49 oil objective heated at 37°C. TIRF analysis was performed using the Roper ILAS module and an Evolve EM-CCD camera (Roper Scientific Instruments). When mentioned, Y-27632 was used at 100 μM. Image acquisition was performed every 12.5 seconds for 30 to 90 minutes.

### Live cell tracking

1205Lu melanoma cells were transfected with IpaA constructs or GFP alone (control) and transferred in microscopy chamber on a 37°C 5%-CO_2_ stage in RPMI1640 medium containing 25 mM HEPES. For cell tracking, samples were analyzed using and inverted Leica DRMIBe microscope and a 20 X phase contrast objective. Image acquisitions were performed every 3 min for 200 hrs. The mean velocity of migration was measured for all tracks followed for at least 5 hours. The root square of MdSD over time was plotted over time and fitted by linear regression. The slopes of the linear fit were compared using an ANCOVA test (linear model). The median cell surface was quantified as the mean of the surface for three time points (25%, 50% and 75%) of the whole cell track and dispersion measured by the Median absolute dispersion (MAD).

### Invasion assays

Tissue culture Transwell inserts (8□μm pore size; Falcon, Franklin Lakes, NJ) were coated for 3□hours with 10□μg of Matrigel following the manufacturer’s instructions (Biocoat, BD Biosciences, San Jose, CA). Inserts were placed into 24-well dishes containing 500□μl of RPMI medium supplemented with 1% fetal calf serum. 5 × 10^4^ melanoma cells were added to the upper chamber in 250□μls of serum-free RPMI medium. After 24 hours, transmigrated cells were scored by bright field microscopy. Experiments were performed at least three times, each with duplicate samples.

### Image processing and statistical analysis

Quantification of vinculin recruitment at the close vicinity of invading bacteria was performed on the sum projection of confocal planes corresponding to *Shigella*-induced actin foci. ROIs were drawn to delineate the actin foci (F) and the bacterial body (b) from the corresponding wavelength channel as shown in Fig. S1. The vinculin recruitment index was calculated as the ratio of the average fluorescence intensity corresponding to vinculin labeling associated with (b) corrected to background over that of (F). For the quantification of the number and size of large adhesion structures induced by *Shigella* invasion in HeLa cells, the confocal fluorescent microscopy plane corresponding to the vinculin-labeled cell basal plane was processed using the imageJ FFT / Bandpass followed by particle analysis plugins with a low size threshold set at 3.54 μm^2^. A semi-automated protocol using Icy software was developed for the quantification of adhesion structures in C2.7 cells (de Chaumont, Dallongeville et al. 2012). Confocal fluorescent microscopy planes were used to detect vinculin structures using HK means thresholding and overlaid binary masks obtained from the threshold projections of F-actin labeled images (Max-entropy method). FAs were detected as spots positive for both vinculin mCherry and actin structures using Wavelet Spot Detector. The number of adhesions was analyzed using Dunn’s multiple comparisons test. The statistical analysis of cell motility was performed in the R software. Medians were compared using a Wilcoxon rank sum test and dispersion by Median absolute dispersion (MAD) parameter.

### Microfluidics cell adhesion assay

Analysis of cell detachment under shear stress was based on previous works (Gutierrez, Petrich et al. 2008). 1205Lu melanocytes were transfected with the indicated constructs, then labeled with 2 μ⍰ calcein-AM (Life Technologies) in serum-free DMEM for 20 minutes. Cells were detached by incubation with 2 μ⍰ Cytochalasin D (Sigma-Aldrich) for 40 minutes to disassemble FAs, followed by incubation in PBS containing 10 mM EDTA for 20 minutes. Cells were washed in EM buffer (120 mM NaCl, 7 mM KCl, 1.8 mM CaCl_2_, 0.8 mM MgCl_2_, 5 mM glucose and 25 mM HEPES at pH 7.3) by centrifugation and resuspended in the same buffer at a density of 1.5 × 10^6^ cells/ml. Calcein-labeled transfected cells and control unlabeled cells were mixed at a 1:1 ratio and perfused onto a 25 mm-diameter glass coverslips (Marienfeld) previously coated with 20 μg/ml fibronectin and blocked with PBS containing 2% BSA (Sigma-Aldrich) in a microfluidic chamber on a microscope stage at 37°C. We used a commercial microfluidic setup (Flow chamber system 1C, Provitro) and a Miniplus3 peristaltic pump (Gilson) to adjust the flow rate in the chamber. Microscopy analysis was performed using a LEICA DMRIBe inverted microscope equipped with a Cascade 512B camera and LED source lights (Roper Instruments), driven by the Metamorph 7.7 software (Universal imaging). Cells were allowed to settle for the indicated time prior to application of a 4 ml/min, flow corresponding to a wall shear stress of 22.2 dyn/cm^2^ (2.22 Pa). Acquisition was performed using a 20 X objective using phase contrast and fluorescence illumination (excitation 480 ± 20 nm, emission 527 ± 30 nm). Fluorescent images were acquired before and after flushing to differentiate between target and control cells. Phase contrast images were acquired every 200 ms. Fold enrichment was defined as the ratio between of attached labeled and unlabeled cells.

## SUPPLEMENTARY INFORMATION

**Supplementary Fig. 1.**
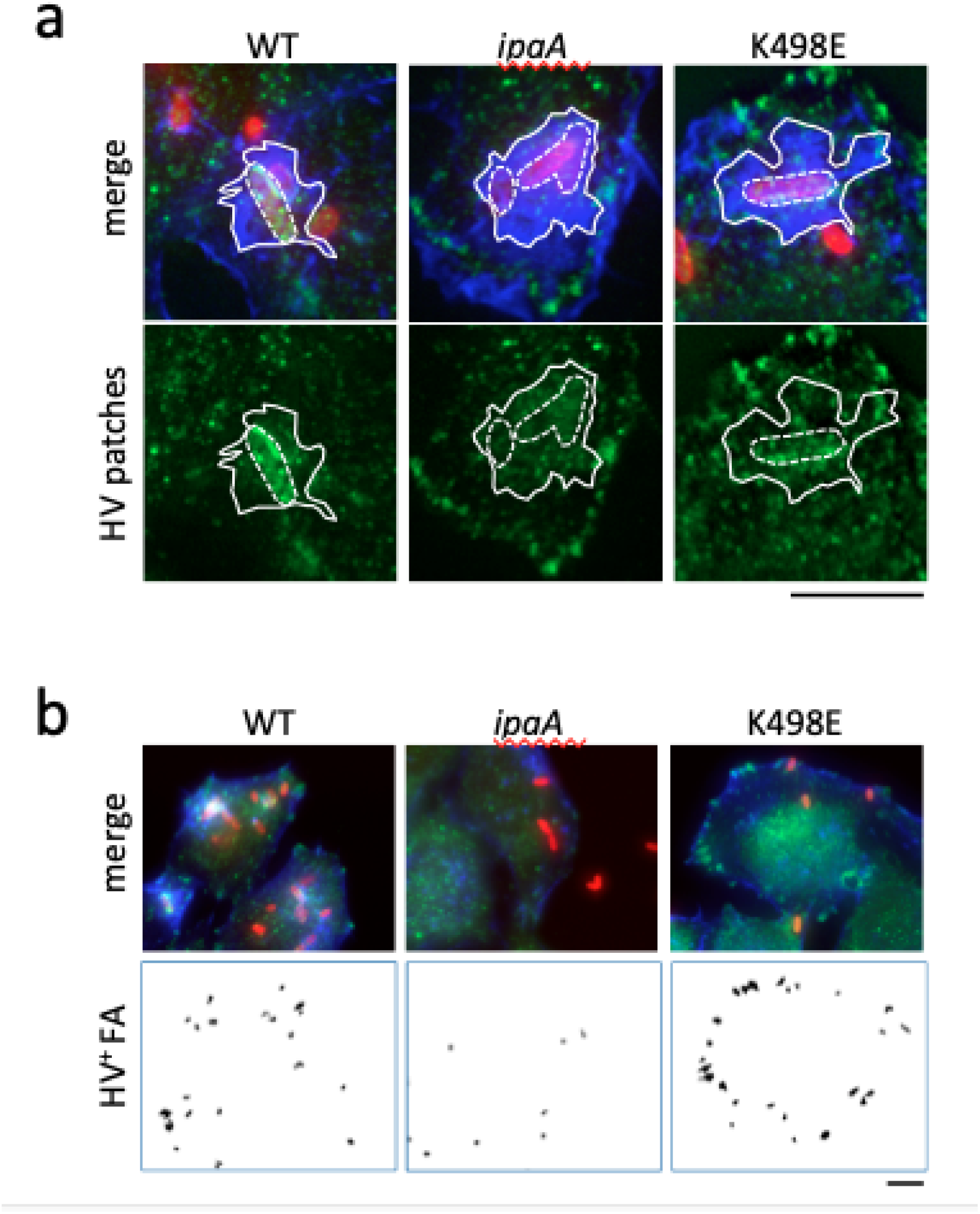
Quantification of IpaA-dependent vinculin recruitment during *Shigella* invasion. HeLa cells were challenged with bacteria for 30 min at 37°C, fixed and processed for immunofluorescence staining. Merge: maximal projections of confocal planes. Red, bacterial LPS; green, vinculin; blue, F-actin. HV: vinculin labeling. **a, b,** representative micrographs. *ipaA* mutant complemented with: full-length IpaA (WT); control vector (*ipaA*); IpaAΔVBS1-2 K498E (K498E). Scale bar = 5 μm. **a,** ROI delineated by: solid lines, actin foci (F); dotted lines, bacterial bodies (b). Vinculin recruitment at the bacterial body was quantified as the ratio of the average fluorescence intensity of vinculin labeling of (b) over that of (F). **b,** HV+ FA: large vinculin adhesions were quantified from images corresponding to the confocal plane of the cell basal surface as described in the Materials and Methods section.

**Supplementary Fig. 2.**
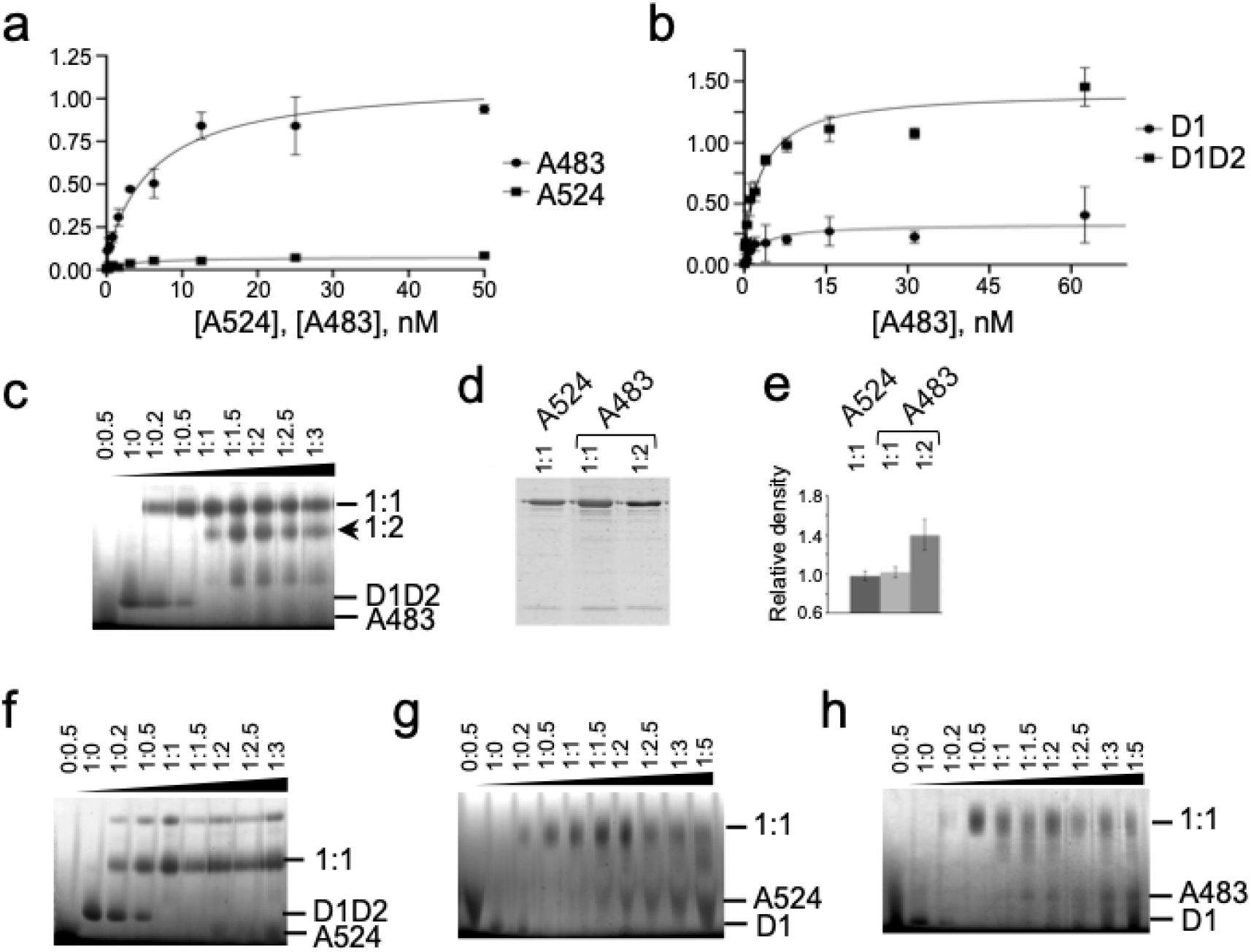
IpaA VBS3 reveals multiple binding sites on vinculin. **a, b,** Solid phase binding assays. **a,** coating: HV; ligands: A483 (solid circles); A524 (solid squares). **b,** coating: D1 (solid circles) or HVD1D2 (solid squares); ligand: A483. **c, f-h,** BN-PAGE in 6-18% polyacrylamide gradient gels and Coomassie staining analysis of D1D2:A483 (**c**), D1D2:A524 (**f**), D1:A524 (**g**) or D1:A483 (**h**) complexes. The molar ratio is indicated above each lane. Arrowheads indicate protein alone, or complex migration at the indicated molar ratio. **d,** bands were recovered from BN-PAGE and analyzed in a second dimension SDS-PAGE in a 15% poly-acrylamide gel and Coomassie staining. Bands were analyzed by densitometry. **e,** ratio of density values for the A524-D1D2 complex (empty bar) and A483-D1D2 complexes corresponding to the upper (light grey bar) or lower (dark grey bar) shifts.

**Supplementary Figure 3.**
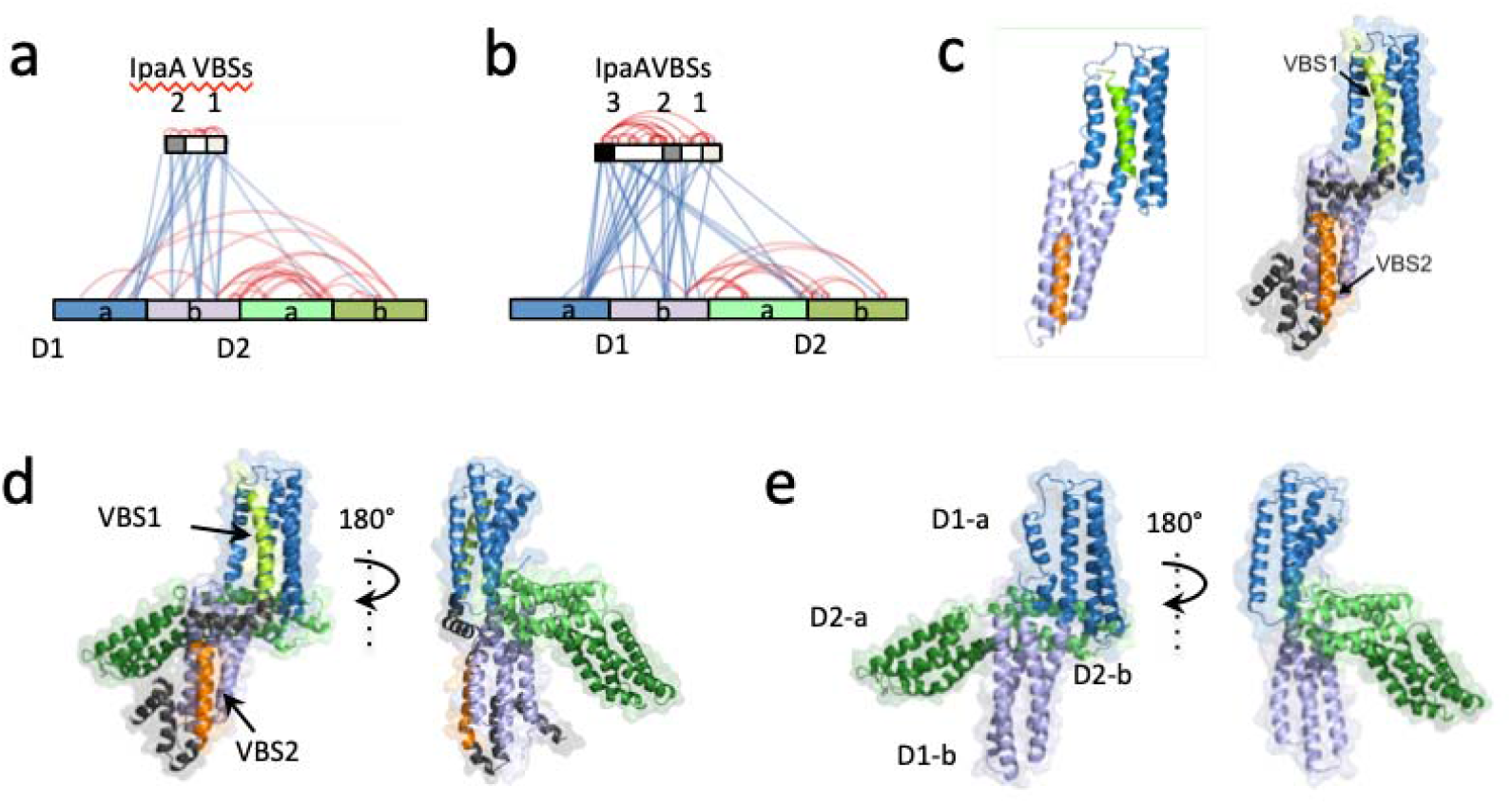
Structural models of vD1:A524. **a, b,** EDC cross-link map from mass spectrometry analysis (LC-MS/MS) of vinculin D1D2-A524 (**a**) and D1D2-A483 (**b**) following extraction of 1:1 complexes from BN-PAGE. Blue lines: inter-molecular links. Red lines: intra-molecular links. Note the links between IpaA VBS3 and the D2 second bundle. Cross-linked residues are detailed in Suppl. Table 1. **c,** left, structure predicted from the resolved vD1: IpaA VBS1: and vD1:IpaA VBS2 crystal structures (Izard, Tran Van Nhieu et al. 2006, Tran Van Nhieu and Izard 2007); right, structural model of vD1:A524. The model was established using RosettaCM protocol and accounts for 19 inter and intra-molecular cross-links out of 24 identified (Suppl. Table 1). Structural models of: **d,** D1D2-A524. **e,** D1D2. IpaA VBS1-2 were docked on the surface of Vinculin D1D2 using MS cross-link constraints. TX-MS protocol in combination with MS constraints was used to unify and adjust the final model, which justifies over 100 cross-links.

**Supplementary Figure 4.**
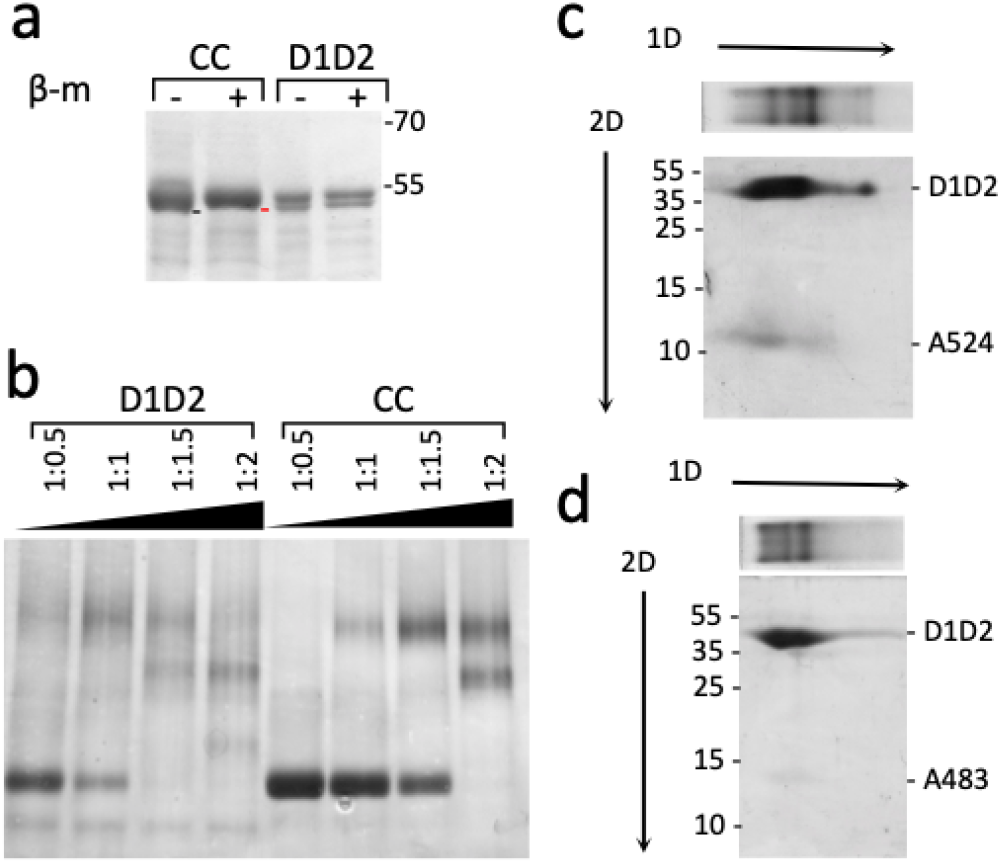
Cysteine-clamped vinculin D1D2 does not form high order complexes. Coomassie blue staining. **a,** Disulfide bridge formation in D1D2. D1D2: wild-type sequence. CC: D1D2 Q68C A396C. SDS-PAGE analysis using a 10 % polyacrylamide gel. + β-metOH: samples were boiled in Laemmli sample loading buffer containing 5 mM beta-mercaptoethanol prior to SDS-PAGE. The molecular weight markers in kDa are indicated. The black and red bars point the respective migration of unreduced and reduced D1D2 Q68C A396C, respectively. **b,** BN-PAGE analysis in a 6-18% polyacrylamide gradient gel of D1D2:A483 (D1D2) and CC:A483 (CC) complexes. The molar ratio is indicated above each lane. Arrowheads indicate protein alone, or complex migration. **c, d,** gel strips from native gels (top) were analyzed in a second dimension as depicted by the 2D arrow by SDS-PAGE using a 10 % polyacrylamide gel. **c,** D1D2 + A524; **d,** D1D2 + A483.

**Supplementary Figure 5.**
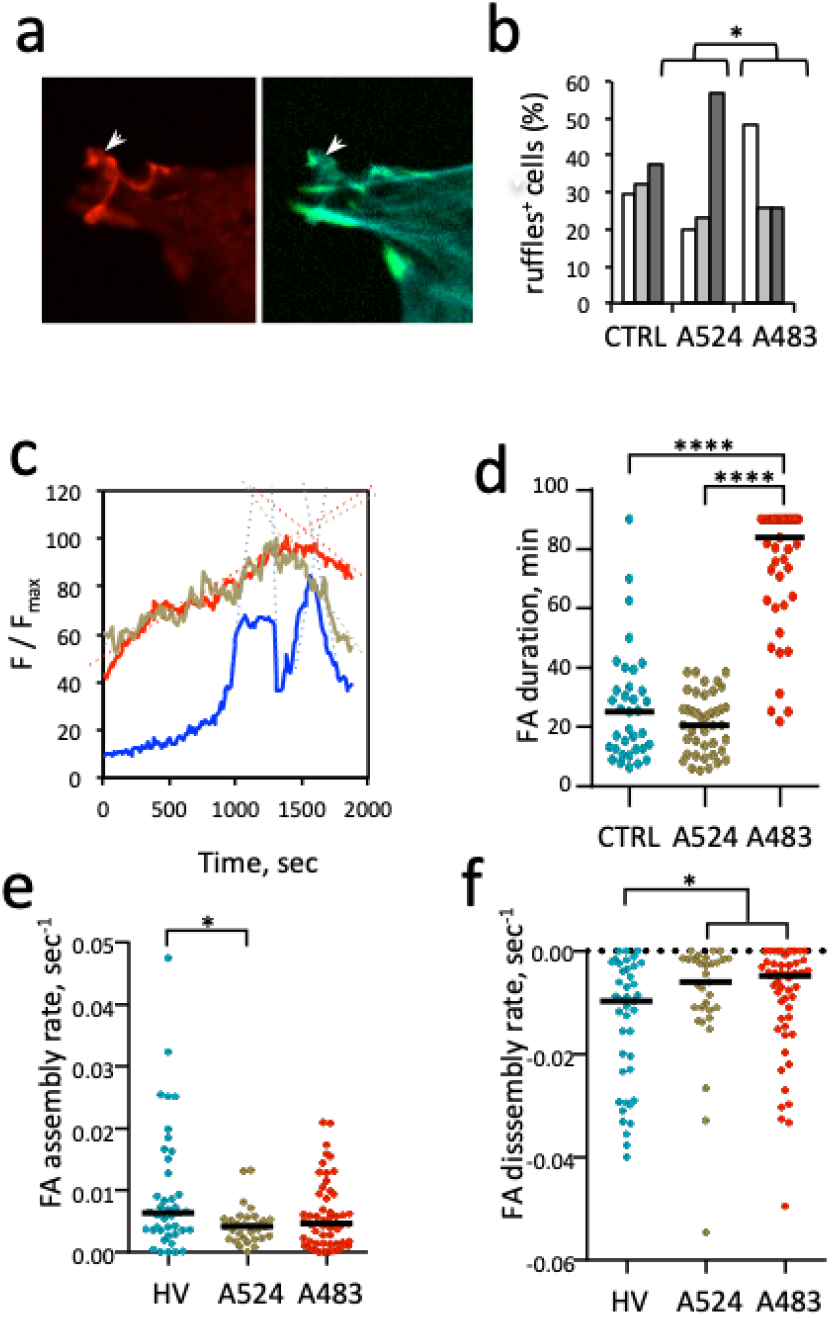
Actin ruffles and TIRF analysis of FA dynamics in A524 and A483 transfectants. **a b,** CTRL: C2.7 cells. A524: GFP-A524 transfectants. A483: GFP-A483 transfectants. **a,** representative fluorescence micrographs. Arrows: adhesions; arrowheads: ruffles. Green: GFP; red: vinculin; cyan: actin. **b,** percent of cells with ruffles ± SEM. Cells with: no ruffles (empty bars); small ruffles (light grey bars); large ruffles (dark grey bars). *: Pearson’s Chi-squared test (N=3, n > 30, p = 0.036). **c-f,** C2. 7 cells were transfected with HV-mCherry (HV), HV-mCherry and GFP-A524 (A524) or HV and GFP-A483 (A483). The dynamics of HV-mCherry-labeled FAs were analyzed by TIRF microscopy. **c,** traces correspond to the variations of average fluorescence intensity of a representative single FA (F) normalized to its maximal average fluorescence intensity over the analyzed period in seconds (F_max_). Blue: HV; green: HV+A524; red: HV+A483. **d,** FA duration. **e, f,** instant assembly (**b**) or disassembly (**c**) rates were inferred from the slopes of linear fits as depicted in a), with a Pearson correlation value > 0.85. HV: n = 41, N = 3; HV+A524: n = 31, N = 2; HV+A483: n = 55, N = 3. Mann-Whitney U test. *: p < 0.05.

## SUPPLEMENTARY TABLES

**Suppl. Table 1.**
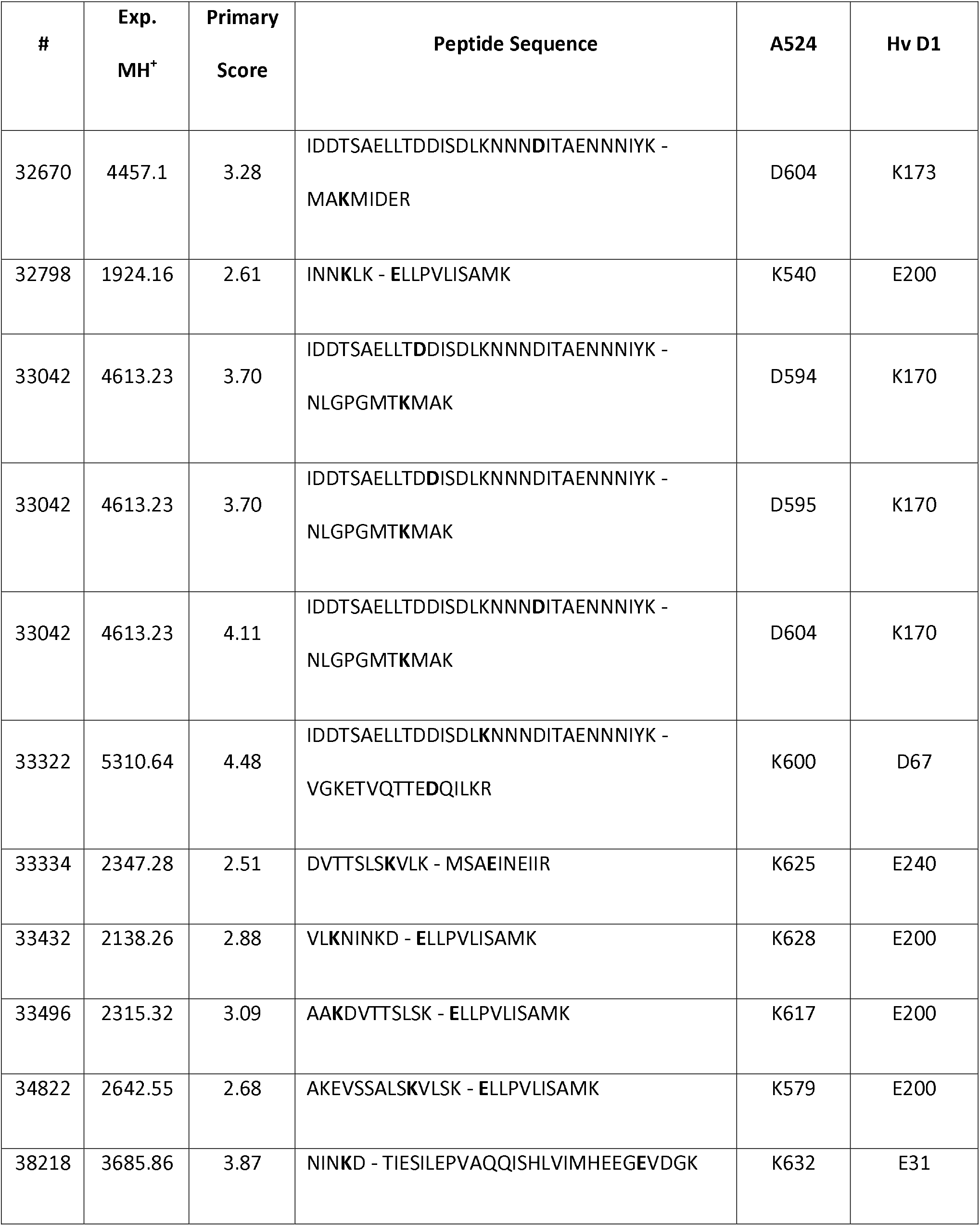
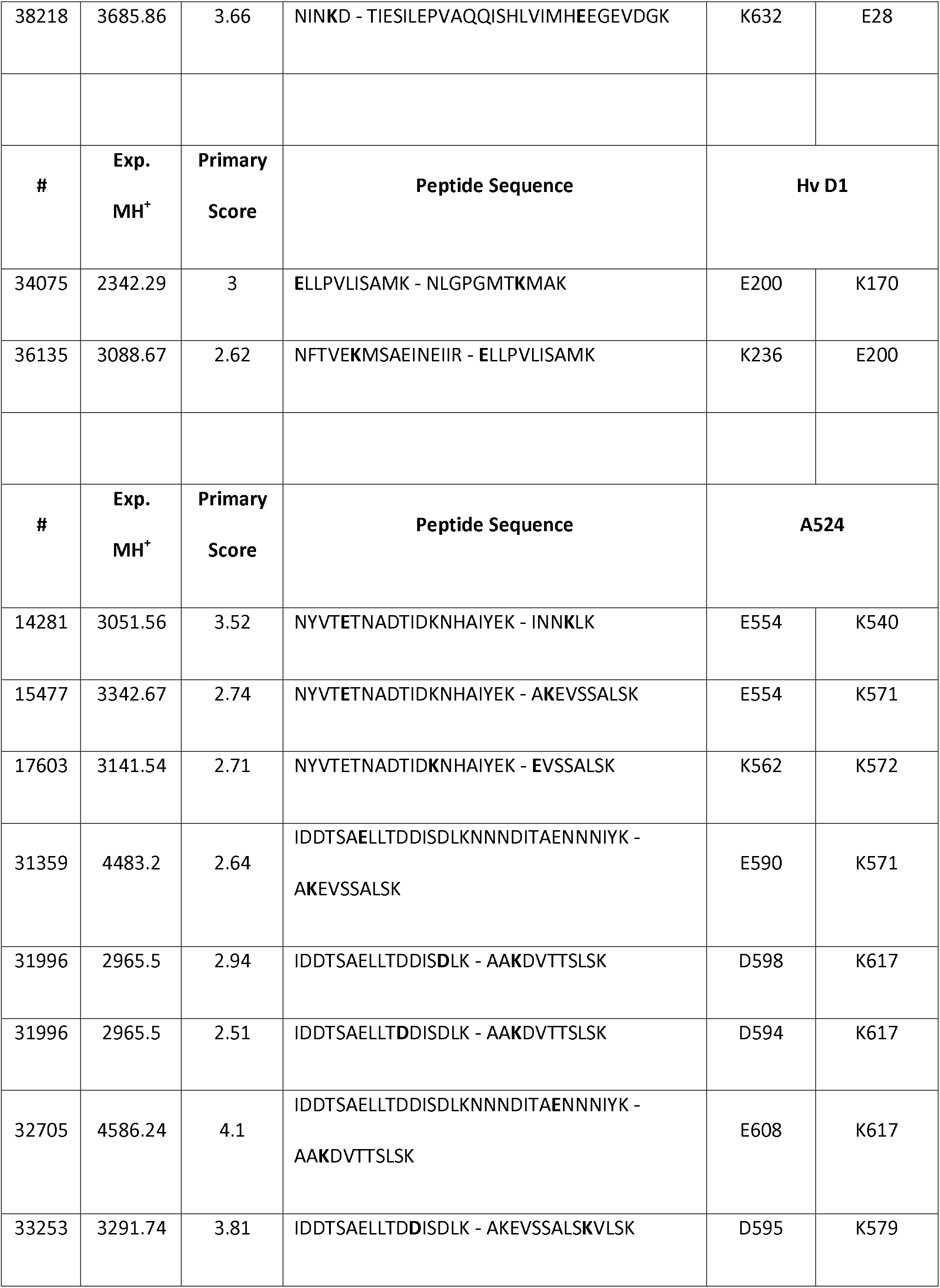
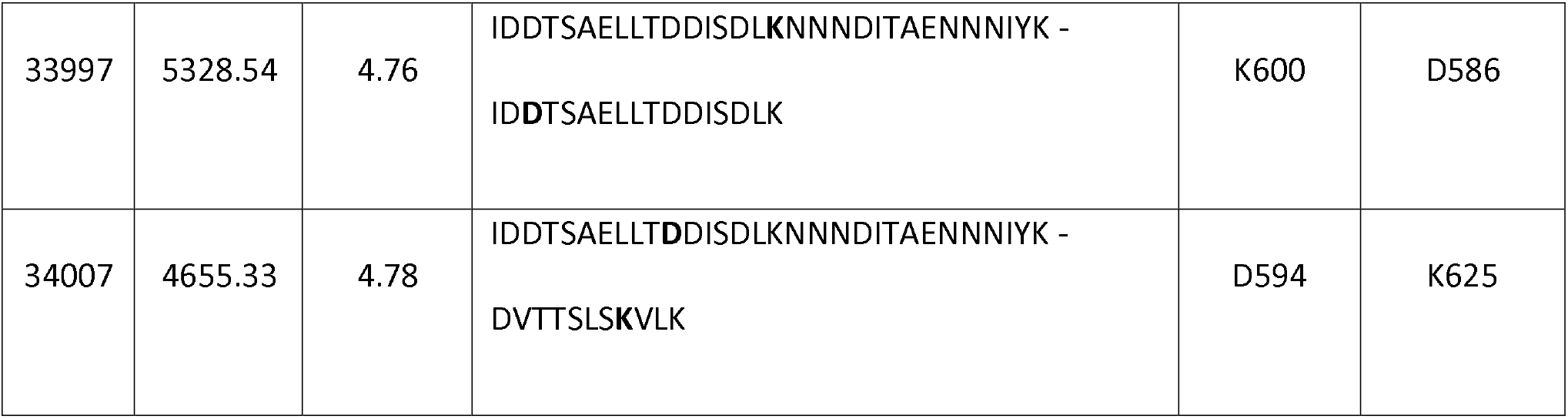
Cross-linked residues characterized in the D1:A524 complex (XL-amino acids of each protein are bolded in the sequences).

**Suppl. Table 2.**
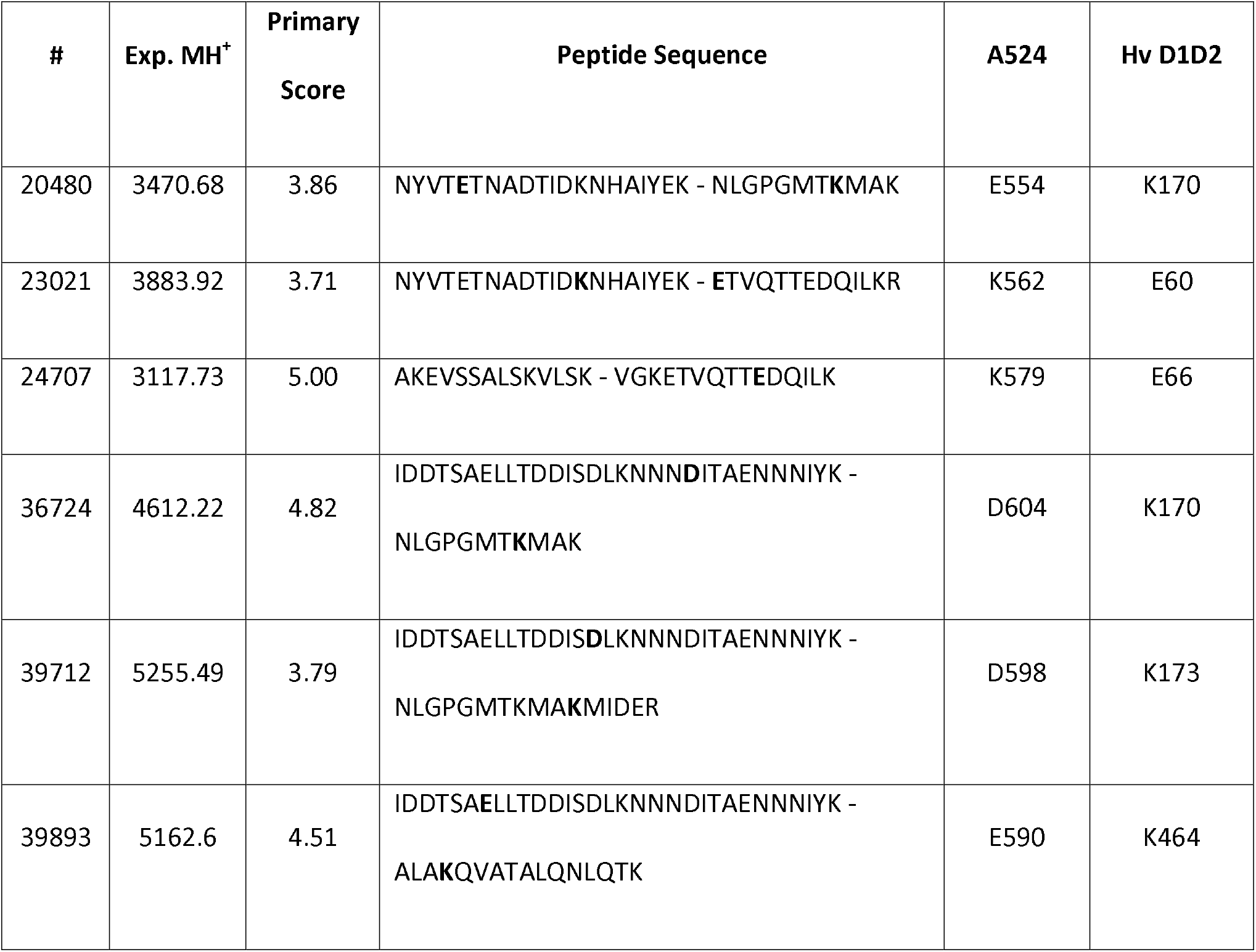
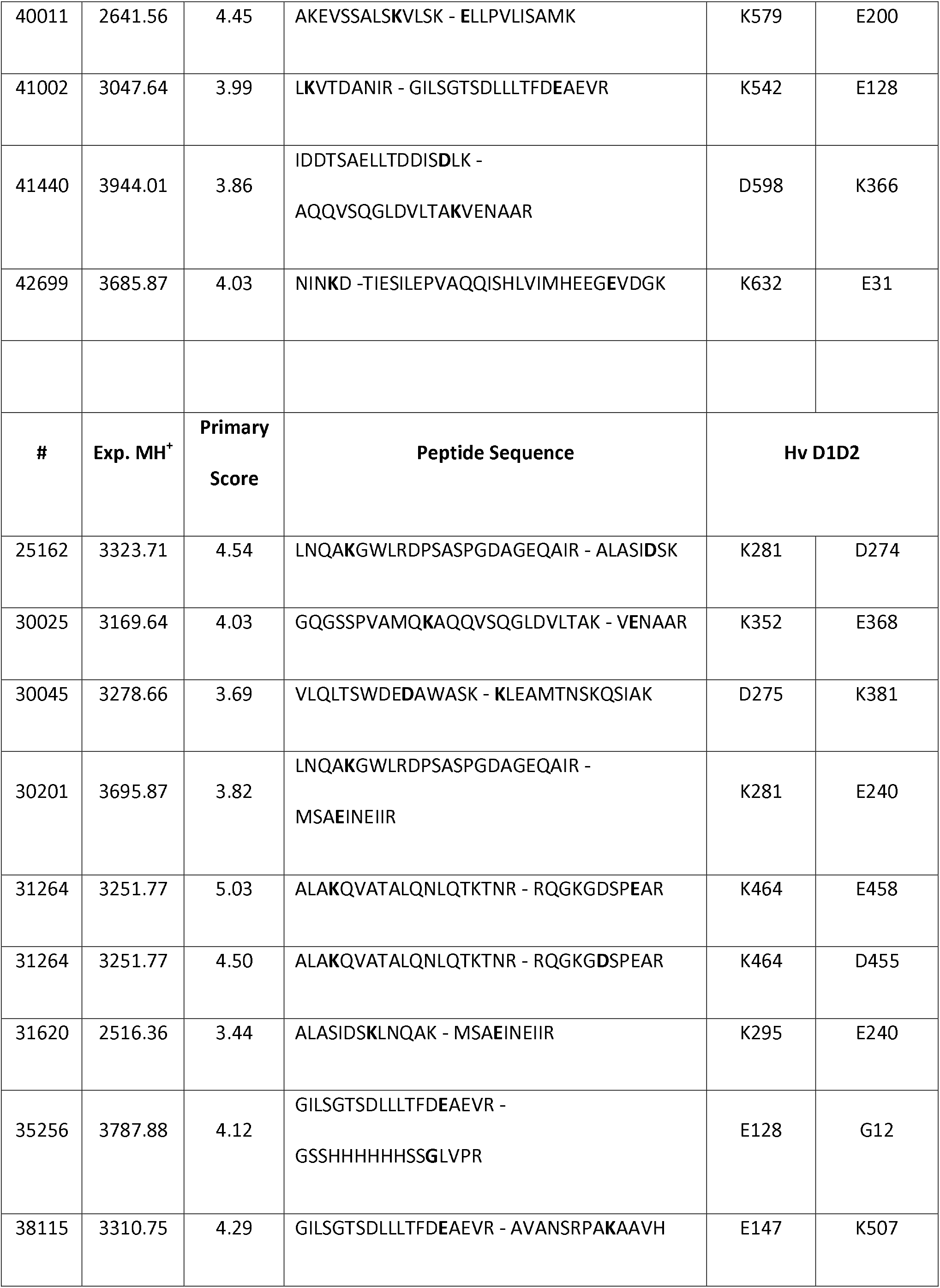
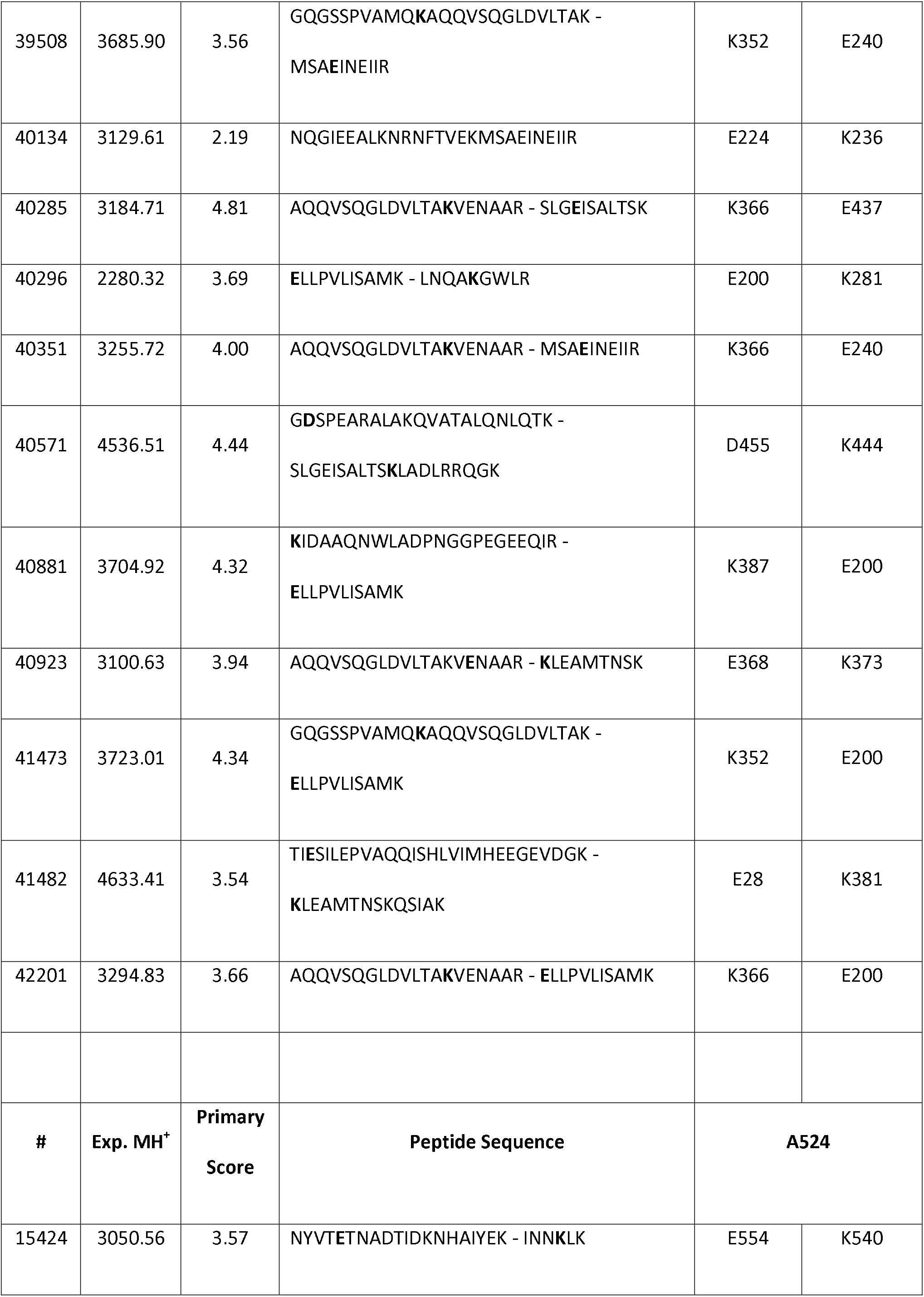
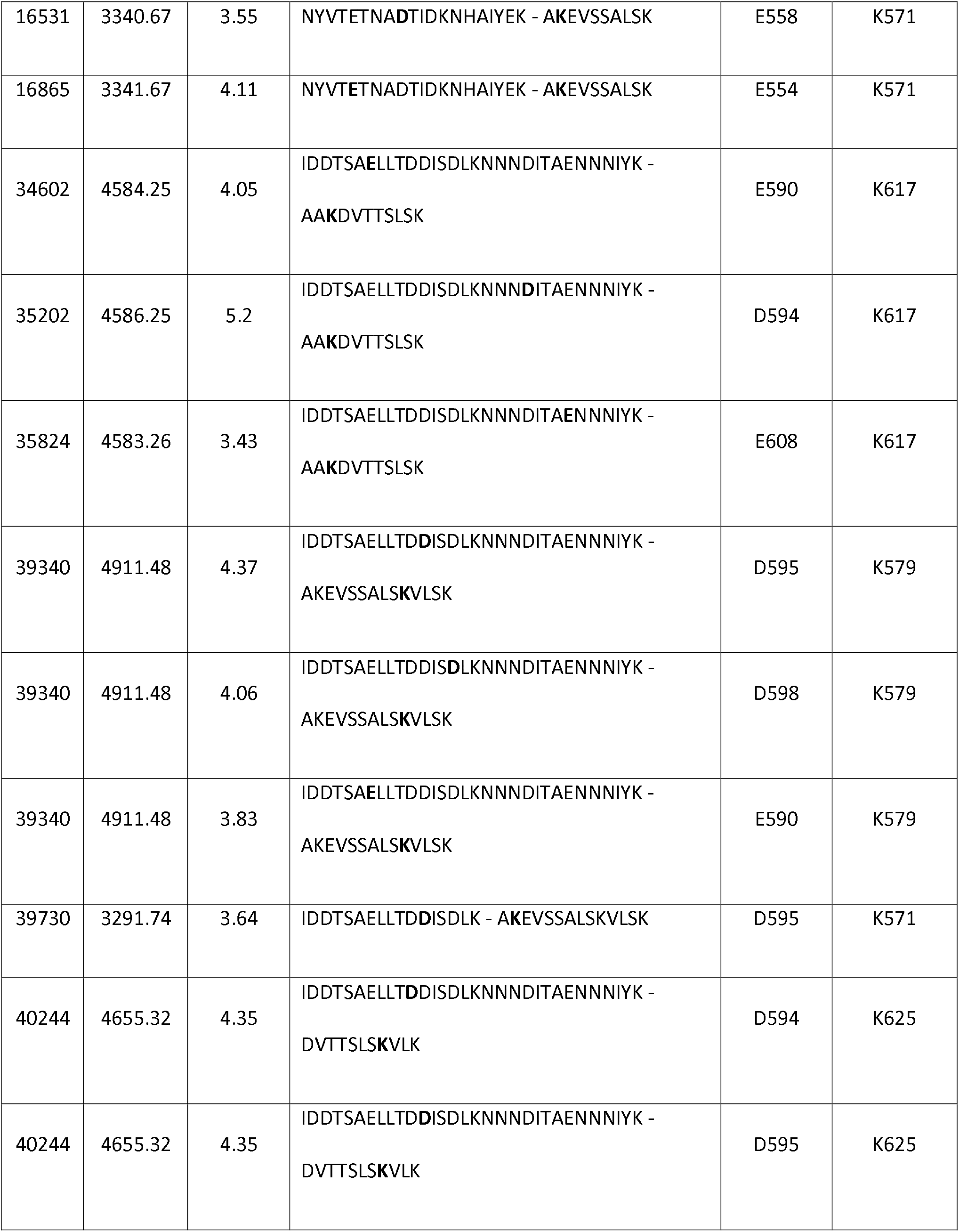
Cross-linked residues characterized in the D1D2:A524 complex (XL-amino acids of each protein are bolded in the sequences).

**Suppl. Table 3.**
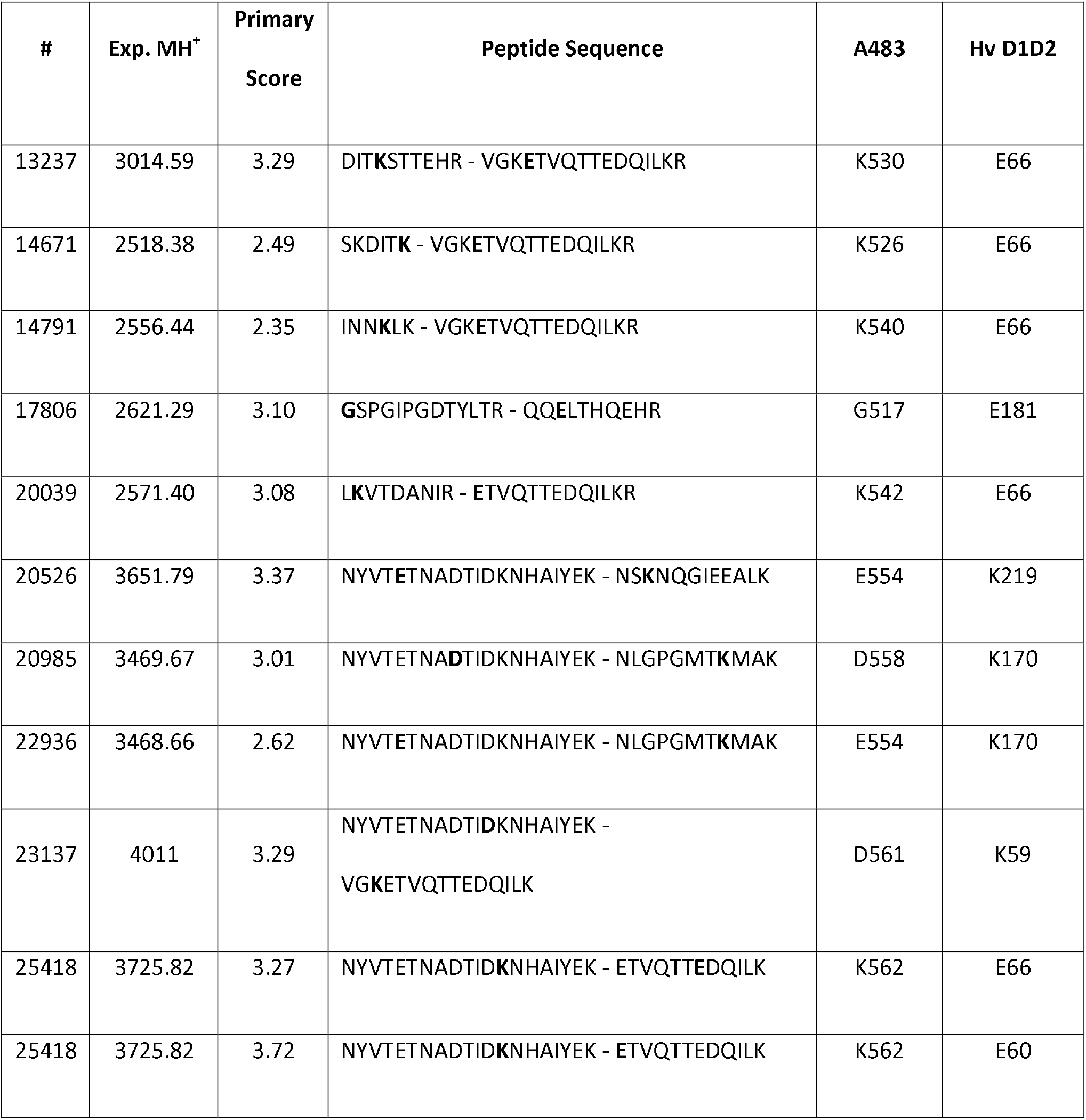
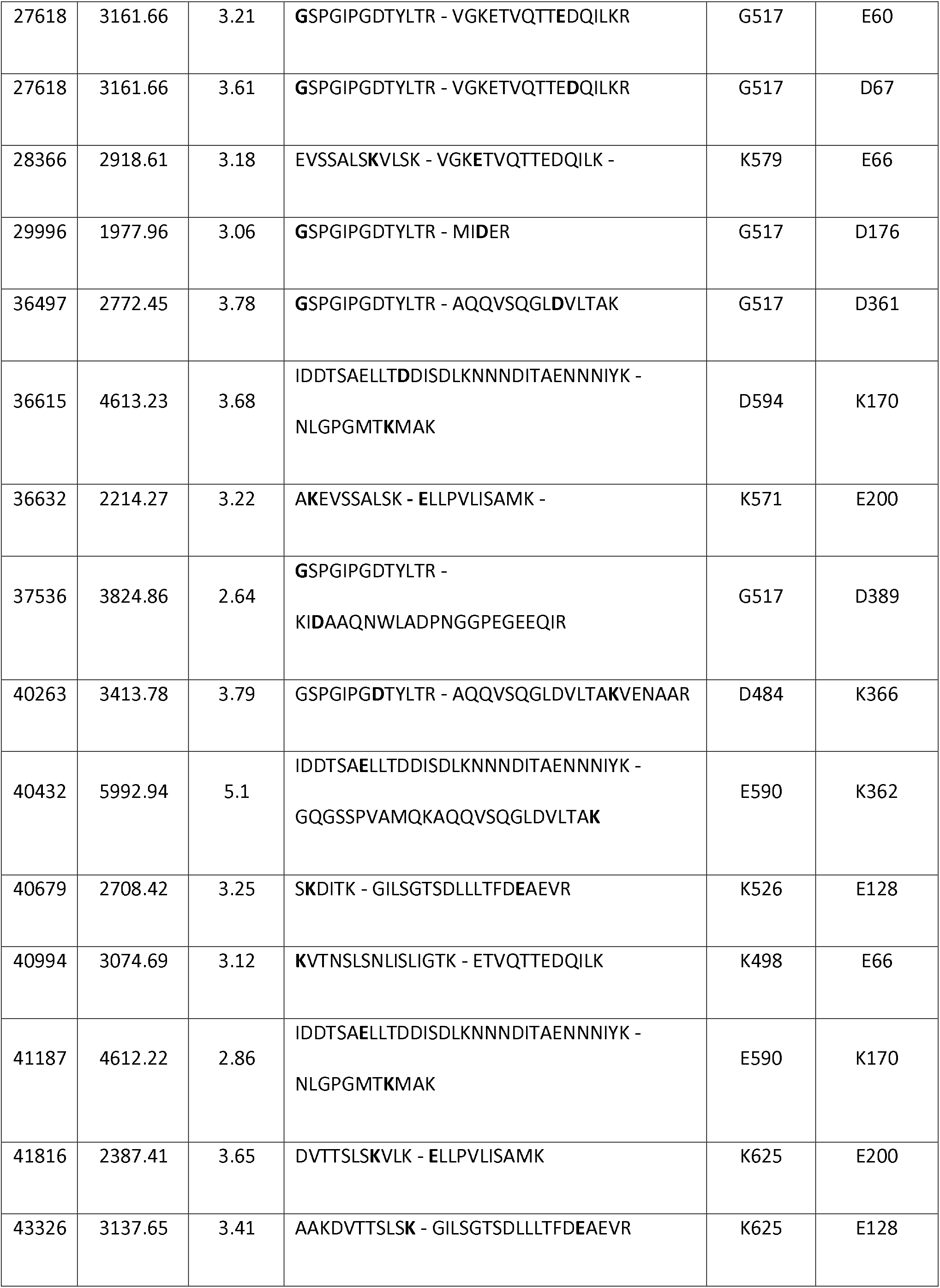
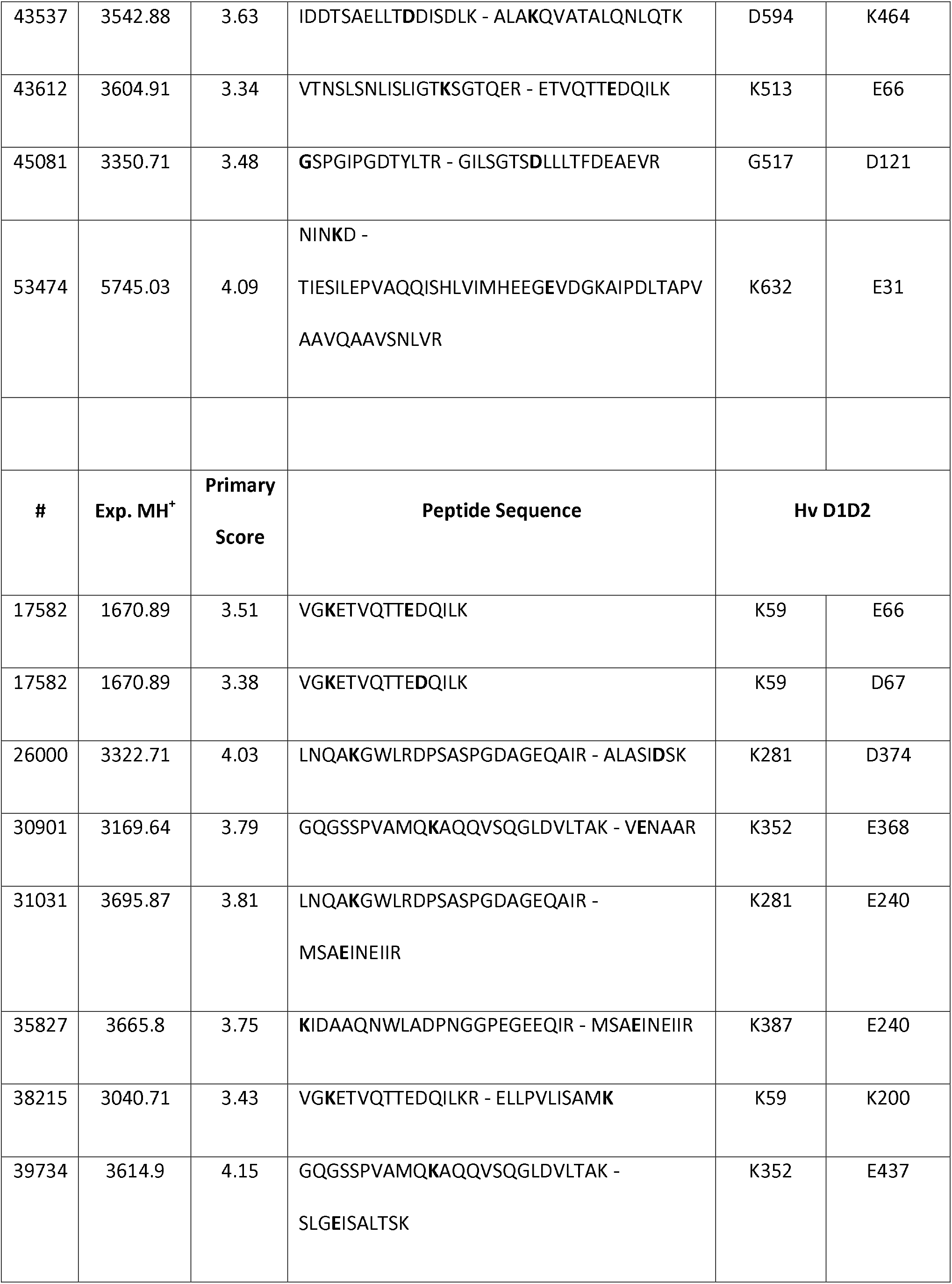
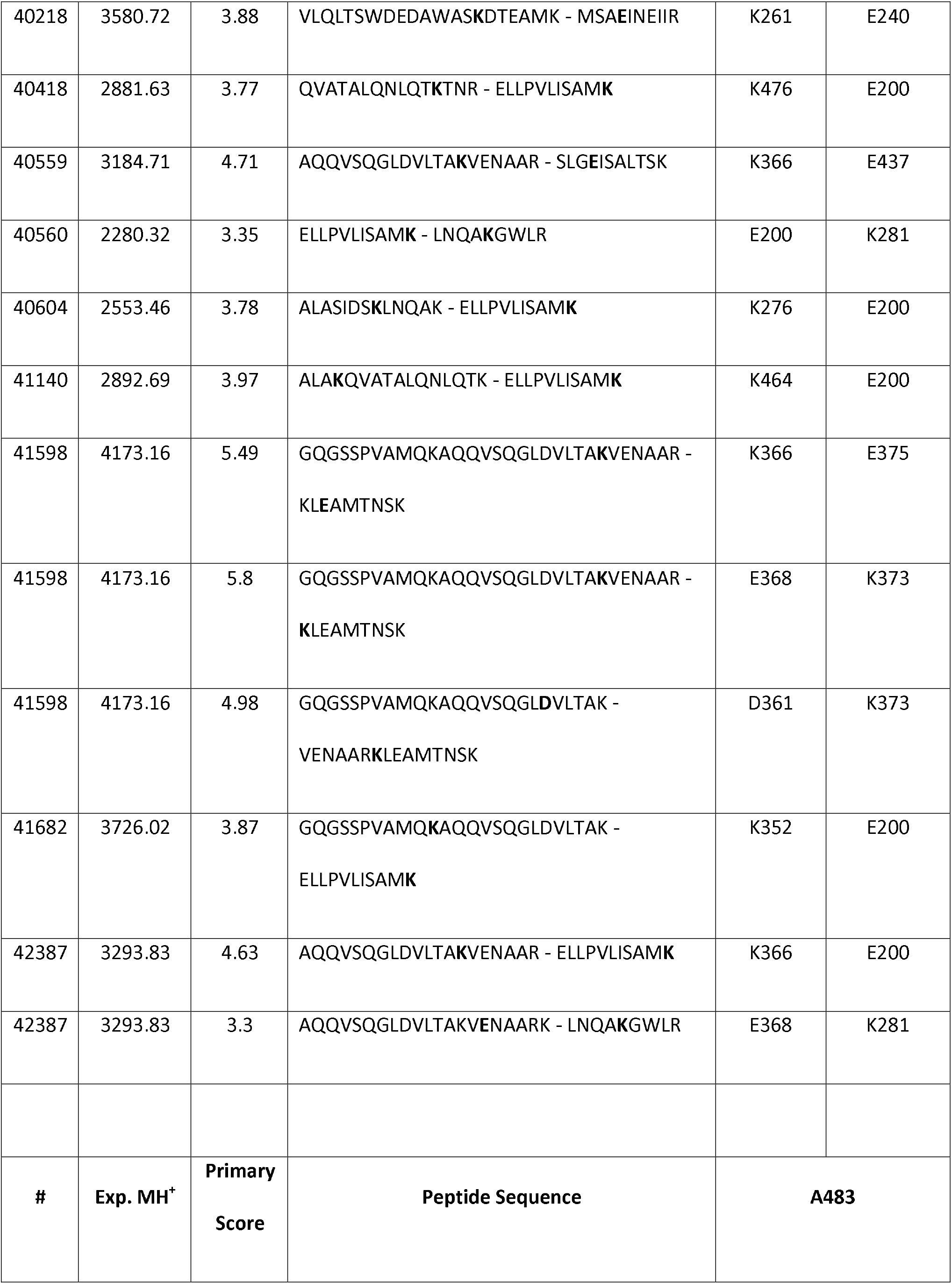
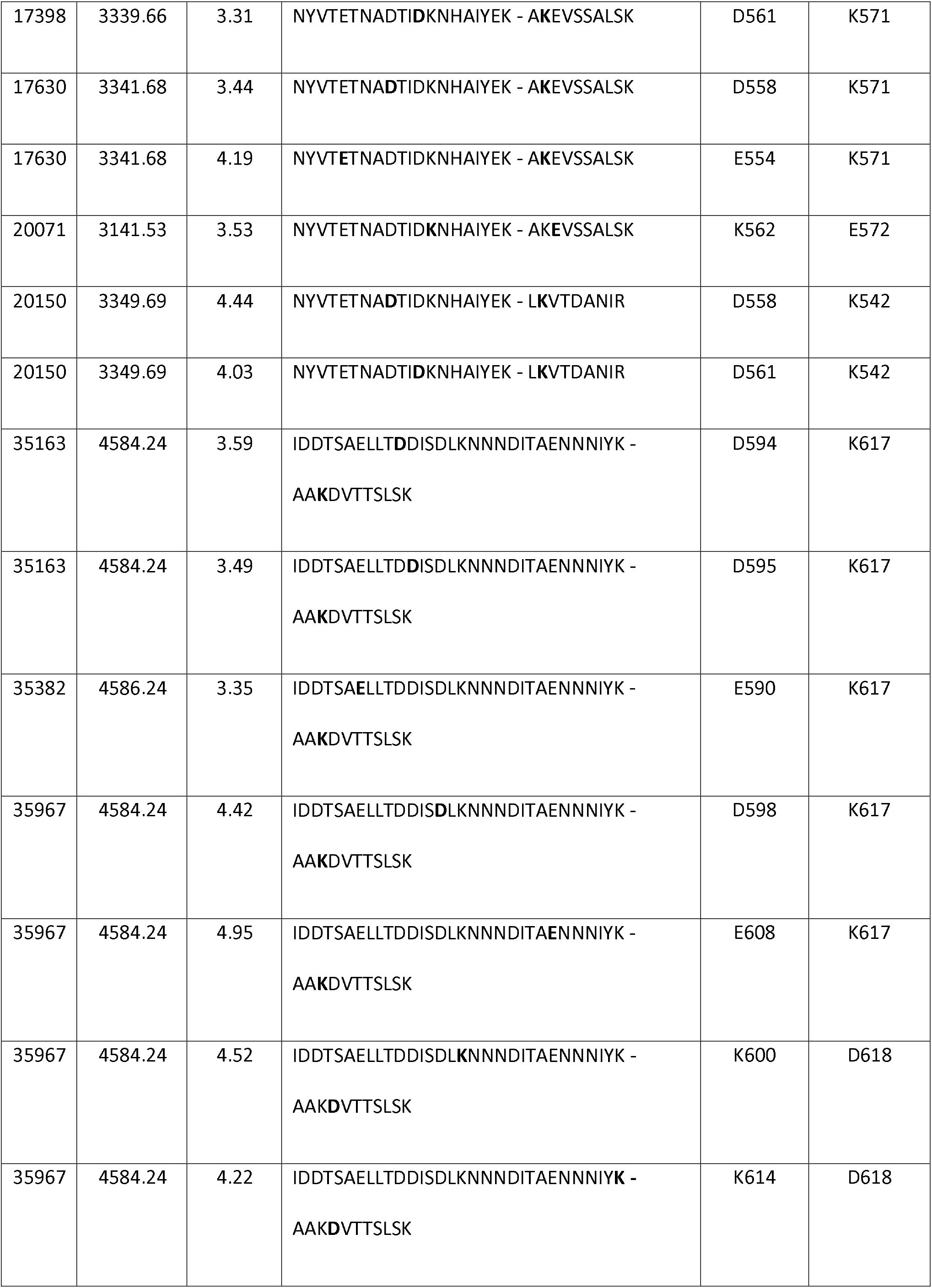
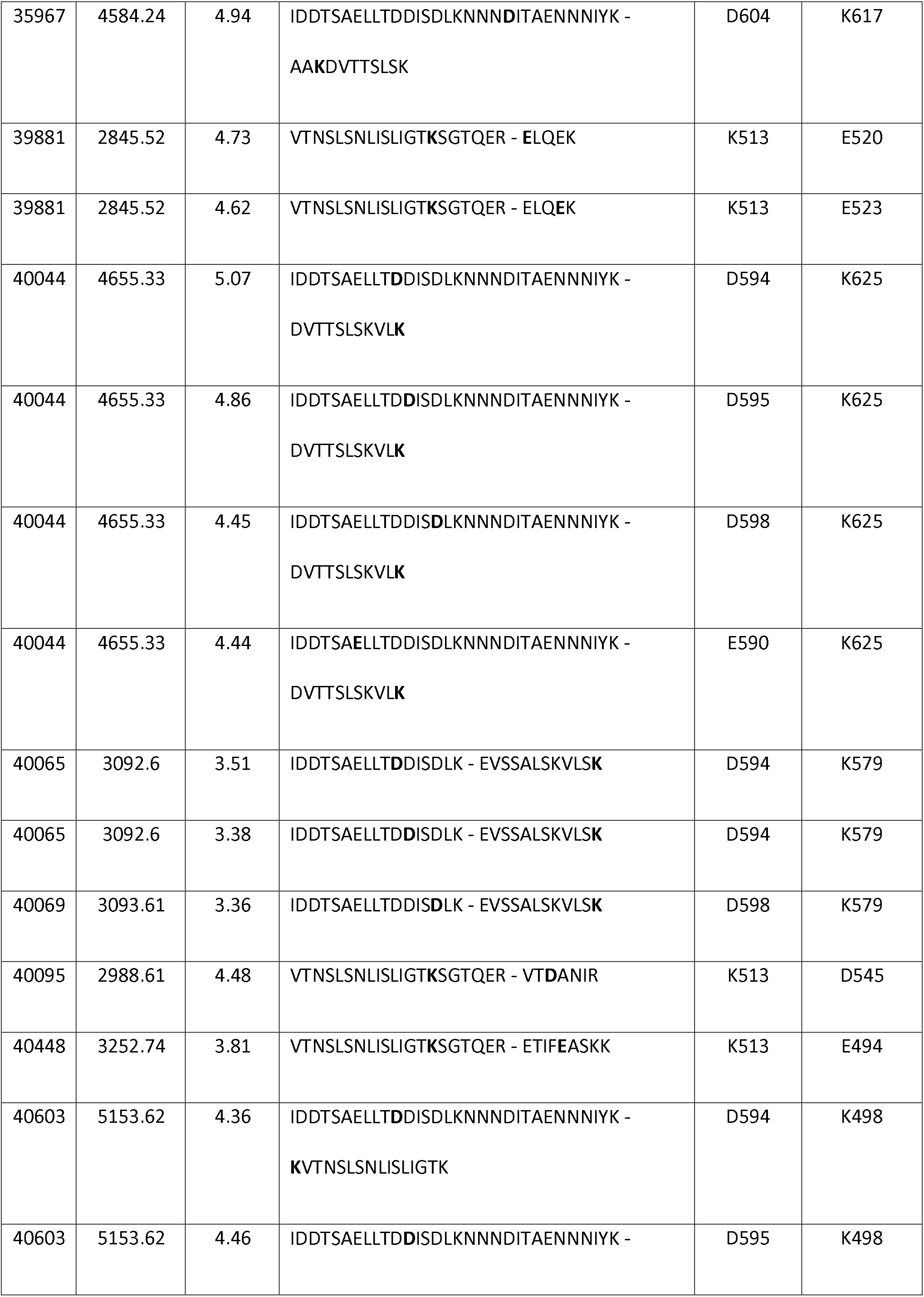
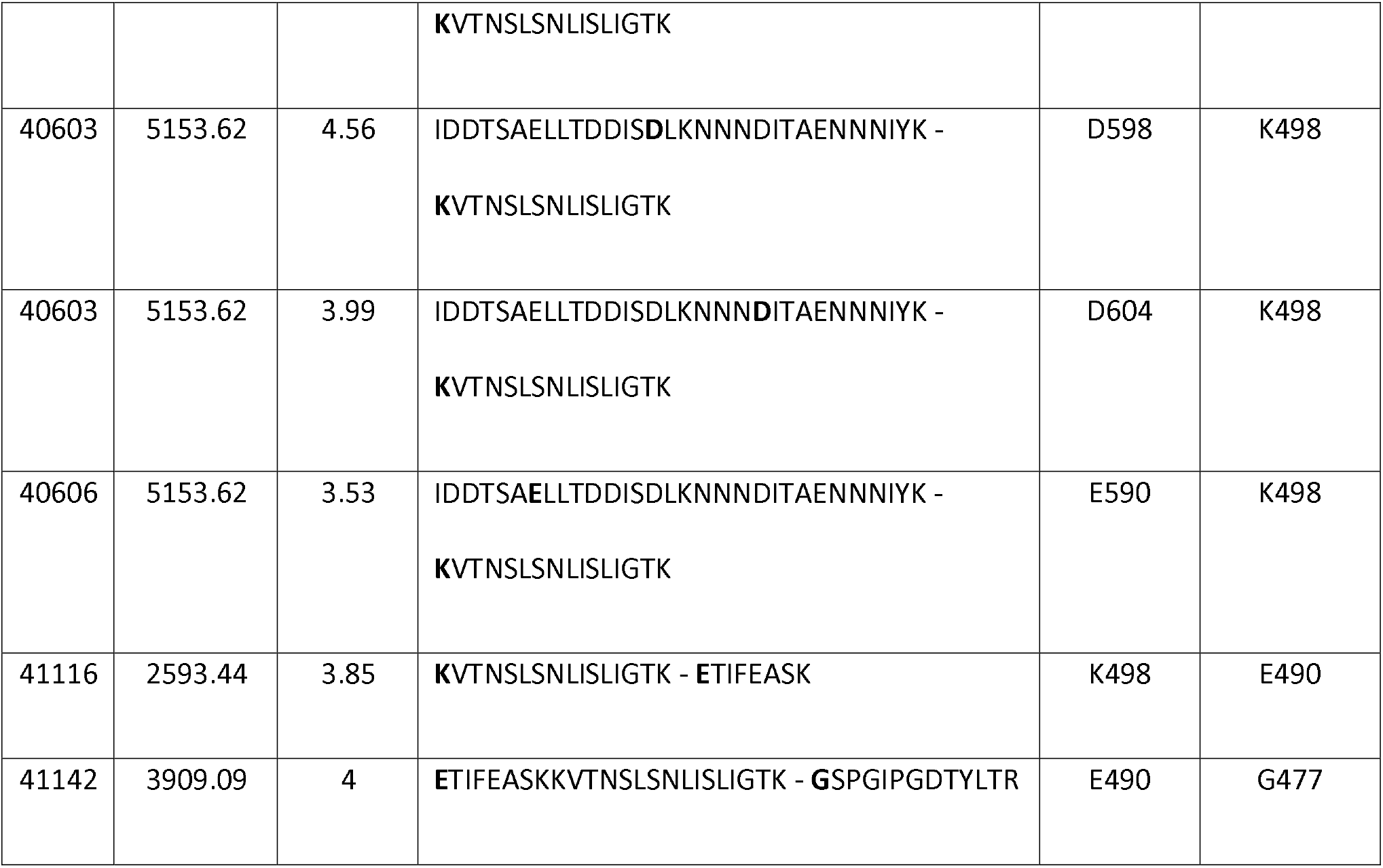
Cross-linked residues characterized in the D1D2 -A483 1:1 complex (XL-amino acids of each protein are bolded in the sequences)

---

**Suppl. movie 1.** TIRF analysis of HV-mCherry expressing C2.7 cells co-transfected with a GFP fusion to the indicated construct. The time is indicated in seconds.

**Suppl. movie 2.** TIRF analysis of HV-mCherry expressing C2.7 cells co-transfected with a GFP fusion to the indicated construct. The time is indicated in seconds. At time “0”, addition of the Rho kinase inhibitor Y-27632 was added at 100 μM final concentration.

**Suppl. movie3.** TIRF analysis of VASP-mCherry expressing C2.7 cells co-transfected with a GFP fusion to the indicated construct. The time is indicated in seconds. At time “0”, addition of the Rho kinase inhibitor Y-27632 was added at 100 μM final concentration.

**Suppl. Movie 4.** 1205Lu melanoma cells 1205Lu melanoma cells were transfected with the indicated constructs. Cells were perfused in a microfluidic chamber and allowed to adhere for 20 min prior to application of shear stress reaching 22.2 dynes.cm^−2^. The elapsed time is indicated in seconds.

